# Extended fractional-polynomial generalizations of diffusion and Fisher-KPP equations on directed networks: Modeling neurodegenerative progression

**DOI:** 10.1101/2023.02.04.527149

**Authors:** Arsalan Rahimabadi, Habib Benali

## Abstract

In a variety of practical applications, there is a need to investigate diffusion or reaction-diffusion processes on complex structures, including brain networks, that can be modeled as weighted undirected and directed graphs. As an instance, the celebrated Fisher-Kolmogorov-Petrovsky-Piskunov (Fisher-KPP) reaction-diffusion equation are becoming increasingly popular for use in graph frameworks by substituting the standard graph Laplacian operator for the continuous one to study the progression of neurodegenerative diseases such as tauopathies including Alzheimer’s disease (AD). However, due to the porous structure of neuronal fibers, the spreading of toxic species can be governed by an anomalous diffusion process rather than a normal one, and if this is the case, the standard graph Laplacian cannot adequately describe the dynamics of the spreading process. To capture such more complicated dynamics, we propose a diffusion equation with a nonlinear Laplacian operator and a generalization of the Fisher-KPP reaction-diffusion equation on undirected and directed networks using extensions of fractional polynomial (FP) functions. A complete analysis is also provided for the extended FP diffusion equation, including existence, uniqueness, and convergence of solutions, as well as stability of equilibria. Moreover, for the extended FP Fisher-KPP reaction-diffusion equation, we derive a family of positively invariant sets allowing us to establish existence, uniqueness, and boundedness of solutions. Finally, we conclude by investigating nonlinear diffusion on a directed one-dimensional lattice and then modeling tauopathy progression in the mouse brain to gain a deeper understanding of the potential applications of the proposed extended FP equations.

## 1. Introduction

A generalization of the Fisher-Kolmogorov-Petrovsky-Piskunov (Fisher-KPP) reaction-diffusion equation [1, 2] to undirected networks has recently been employed to study the spreading of prion-like proteins within the brain [3–6]. In fact, the Fisher-KPP equation has been altered by replacing the continuous Laplacian operator with the conventional graph Laplacian operator. This generalization and the concept of nonlinear diffusion prompted us to propose a new nonlinear graph Laplacian operator that allows us to capture more complex diffusion phenomena on both undirected and directed networks. We also extend the reaction term of the Fisher-KPP equation to introduce a larger family of reaction-diffusion models.

The dynamics of linear Laplacian operators in directed graphs have been well studied [7–10]. In the literature, the standard graph Laplacian has been developed in a variety of ways to study spectral information, such as eigenvalues, eigenvectors, and Cheeger constants, for graphs and hypergraphs [11–18]. Aside from the extensions regarding spectral graph theory, several generalizations of the standard graph Laplacian exist particularly for exploring anomalous and nonlinear diffusion processes, most of which are restricted to undirected graphs [19–31], and for studying the dynamics of chemical reaction networks (CRNs) [32], which are directed networks [33, 34]. Fractional graph Laplacian operators constructed by raising the Laplacian matrix to a real power between 0 and 1 [19–22], d-path Laplacian operators defined by using path matrices accounting for the existence of shortest paths of length d between two nodes [23], and the Mellin-transformed d-path Laplacian operators 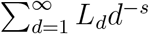 [24–26], where *s* is a nonnegative real number, have been applied to investigate superdiffusion with a linear differential equation. Besides the normal diffusion, in which the mean square displacement (*MSD*) of particles scales linearly with time, and the anomalous superdiffusion processes, in which *MSD* (*t*) is proportional to *t^ζ^* with *ζ* > 1, there is a substantial body of knowledge about the subdiffusion processes for which *ζ* < 1, especially observed in biological systems [35–41]. In a recent paper [27], Diaz-Diaz and Estrada presented a linear diffusion equation incorporating the Mellin-transformed d-path Laplacian operator, which utilizes fractional-time derivatives to describe subdiffusion on undirected networks. Some nonlinear generalizations of the standard graph Laplacian operator on undirected graphs are the p-Laplace operator 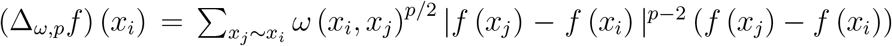 for a vertex *x_i_* and its variants, which have been exploited to introduce a family of discrete-in-time diffusion equations used in image processing [28, 29]. In order to examine the dynamics of interacting random walkers moving over unweighted undirected graphs, a nonlinear graph Laplacian operator has been suggested as follows [30]: 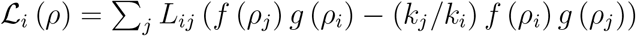 for the mean-field node density *ρ_i_*, where *L_ij_* = *a_ij_/k_j_* – *δ_ij_* is the usual graph Laplacian and *k_i_* represents the degree of node *i*. Given functions *f* (*x*) = *x^α^* and *g* (*x*) = (1 – *x*)^*σ*^ for 0 ≤ *x* ≤ 1 and zero elsewhere, it was illustrated that the diffusion equation formed by 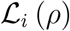 has a unique stationary solution under some assumptions. Nevertheless, the authors have not addressed the existence and uniqueness of solutions of the proposed diffusion equation in a broader sense. In a recent work [31], considering functions *f* (*x*) = *x^m^* and *g* (*x*) = 1, it has been shown that taking the continuum limit of the diffusion equation incorporating 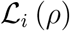 for an infinite q-homogeneous tree, where each node has degree *q* + 1 and the distance between nodes is given by *a*, yields the nonlinear partial differential equation (PDE) *∂_tρ_* = *a*^2^ (*q* + 1) *∂_xx_* (*ρ^m^*) − *a* (*q* − 1) *∂_x_* (*ρ^m^*), where *ρ* (*x, t*) is identified with *ρ* (*x, t*) = *ρ_i_* (*t*). In the case of an infinite regular lattice, i.e., *q* = 1, the aforementioned PDE reduces to the so-called porous medium equation *∂_t_ρ* = *σ*Δ (*ρ^m^*), with *m* > 1, admitting a family of self-similar solutions (in a weak sense), called Barenblatt-Pattle solutions, representing diffusion from a point source [42, 43]. The article [31] also verified the uniqueness of the stationary solution under specific conditions. However, similar to [30], it did not discuss the existence and uniqueness of the solution of the graph diffusion equation 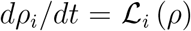 with initial conditions *ρ_i_* (0) ≥ 0 in a more general sense. Recently, Veerman et al. [32] propounded a nonlinear Laplacian framework for CRNs, which leads to the polynomial system of differential equations 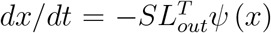, where *S* is a non-negative matrix with no zero rows, *L_out_* is an out-degree Laplacian matrix, and each element of vector *ψ* is a monomial. It has been demonstrated that for a componentwise strongly connected network when *KerS* ∩ *ImL^T^* = {0}, the proposed polynomial system has exactly one positive equilibrium *x\** in a specified invariant set *X_z_*. Using a Lyapunov function, the authors concluded the (local) asymptotic stability of that equilibrium point *x\** in *X_z_*, and they also verified that the *ω*-limit set of any positive initial condition in *X_z_* either equals *x\** or is a bounded set contained in the boundary of the positive orthant.

Our generalizations of the diffusion and reaction terms result in systems of differential equations involving fractional polynomial (FP) functions whose definitions are restricted to the nonnegative orthant, which is a closed subset of the Euclidean space. Hence, we first extend these functions to functions on an open subset possessing a differentiable structure, empowering us to gain the common methods and theorems of dynamical systems theory to analyze the proposed FP equations. In addition, since the extended FP functions are continuously differentiable on the whole space, we can investigate the properties of solutions not only with nonnegative initial conditions but also negative ones whose practical importance can be realized by introducing the notion of active concentration. The elaboration of the details is continued in Subsection 3. The remainder of the paper is organized as follows; we propose a nonlinear Laplacian operator and its corresponding diffusion equation in Subsections 4.1 and 4.2, respectively. For the extended FP diffusion equation, the existence and uniqueness of solutions as well as positively invariant sets are discussed Subsubsection 4.2.1, and we also establish the convergence of solutions and stability of equilibria in Subsubsection 4.2.2. Next, our generalization of the Fisher-KPP equation and a family of its positively invariant sets are presented in Subsection 5. For the purpose of illustrating the potential applications of our model, we first examine nonlinear diffusion on a directed one-dimensional lattice in Section 6 and then proceed to model tauopathy progression in the mouse brain in Section 7.

## 2. Mathematical preliminaries

### 2.1. Definitions and notations

To delineate our model, we first need to introduce a few definitions and notations. The set of all real numbers will be denoted by ℝ, and given *γ* ∈ ℝ, we define ℝ_*≥γ*_ = [*γ*, ∞), ℝ_>*γ*_ = (*γ*, ∞), ℝ_*≤γ*_ = (−∞, *γ*], and ℝ_*<γ*_ = (–∞, *γ*). Accordingly, 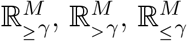, and 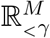 represent the sets of all *M*-tuples whose components belong to ℝ, ℝ_*≥γ*_, ℝ_>*γ*_, ℝ_*≤γ*_, and ℝ_*<γ*_, respectively. The set of all *M*-by-*M* matrices over ℝ is indicated by 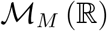, and ***I*** also denotes the identity matrix. Throughout the paper, boldface is used to distinguish vectors and matrices from scalars. Suppose ***u***, ***v***, ***w*** ∈ ℝ^*M*^, and let **Λ** (***v***) be a diagonal matrix whose diagonal elements are the elements of ***v***. Then, the vector ***v*** ⊙ ***w*** defined by ***v*** ⊙ ***w*** = **Λ** (***v***) ***w*** is called the Hadamard product of ***v*** and ***w***, which is commutative, i.e., ***v*** ⊙ ***w*** = ***w*** ⊙ ***v***, associative, i.e., ***u*** ⊙ (***v*** ⊙ ***w***) = (***u*** ⊙ ***v***) ⊙ ***w***, and distributive over vector addition, i.e., ***u*** ⊙ (***v*** + ***w***) = ***u*** ⊙ ***v*** + ***u*** ⊙ ***w*** [44]. We also define functions 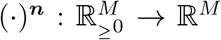 by 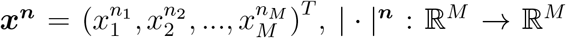 by 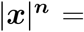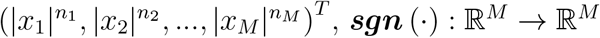 by ***sgn*** (***x***) = (*sgn*(*x*_1_), *sgn*(*x*_2_), *…, sgn*(*x_M_*))^*T*^, and ***ln*** (| · |) : ℝ^*M*^ → ℝ^*M*^ by 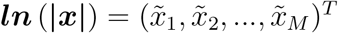 where

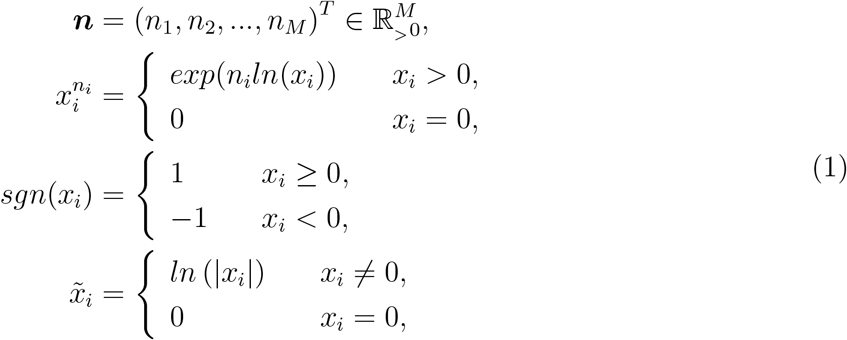

and |*x_i_*| is the absolute value of *x_i_*. For notational simplicity, we will use ***x**^n^* and |***x***| rather than 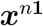 and |***x***|^**1**^, respectively, where **1** is an all-ones vector with an appropriate dimension. It is also obvious that ***x*** = **Λ** (***sgn***(***x***)) |***x***| and |***x***|^***n***^ = **Λ** (|***x***|^***n***^) **1**.

### 2.2. Autonomous systems of ODEs

Now, we recall some theorems and results concerning autonomous systems of differential equations, which will be used in our analysis. Consider the system

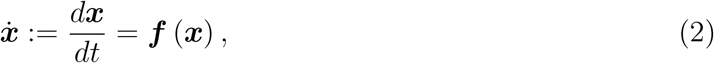

where ***f*** : *X* → ℝ^*M*^ is a continuously differentiable function from a domain (open and connected set) *X* ⊆ ℝ^*M*^ into ℝ^*M*^. Then, a point ***x**_e_* is said to be an equilibrium point of (2) if ***f*** (***x**_e_*) = 0.

#### Theorem 2.1.

*[45] Let* Ξ *be a compact (closed and bounded) subset of X, and **x***_0_ ∈ Ξ. *If every solution of the system (2) with the initial condition **x***(0) = ***x***_0_ *lies entirely in* Ξ, *then there is a unique solution that is defined for all t* ≥ 0.

#### Theorem 2.2

(Lyapunov’s stability theorem). *[45] Suppose that V* : *X\** → ℝ *is a continuously differentiable function defined in a domain X\** ⊆ *X which contains an equilibrium point **x**_e_ of the system (2), and let* ∇*V* (***x***) *be its gradient. If V* (***x***) *and its derivative along the trajectories of the system (2), i.e.*, 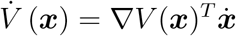, *satisfy the following conditions*

- *V* (***x***) *is positive definite with respect to **x**_e_; and*
- 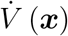 is negative semidefinite.

*Then **x**_e_ is stable in the sense of Lyapunov, and V* (***x***) *is called a Lyapunov function*.

We say that *V* (***x***) is positive (resp., negative) definite with respect to ***x**_e_* if *V* (***x**_e_*) = 0 and *V* (***x***) > 0 (resp., −*V* (***x***) > 0) for all ***x*** ∈ *X\** \ {***x**_e_*}. Further, if it satisfies the weaker condition *V* (***x***) ≥ 0 (resp., −*V* (***x***) ≥ 0), it is said to be positive (resp., negative) semidefinite. It is not difficult to check the sign definiteness of a quadratic function ***x**^T^ **P x*** where ***P*** is a real symmetric matrix. It can be shown that ***x**^T^ **P x*** > 0 (resp., ***x**^T^ **P x*** ≥ 0) for all ***x*** ≠ 0 if and only if every eigenvalue of ***P*** is positive (resp., nonnegative), in which case the matrix ***P*** is called positive definite (resp., positive semidefinite) and denoted by ***P*** ≻ 0 (resp., ***P*** ⪰ 0).

#### Theorem 2.3

(LaSalle’s invariance theorem). *[46] Let* Ξ^*\**^ ⊂ *X be a compact set that is positively invariant with respect to the system (2), i.e*.,

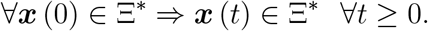

*If an equilibrium point **x**_e_ belongs to* Ξ^*\**^ *and there exists a continuously differentiable function V* : *X* → ℝ *such that*

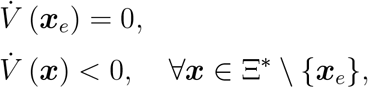

*then every solution starting in* Ξ^*\**^ *approaches **x**_e_ as t goes to infinity*.

Note that Theorem 2.3 is actually a corollary of LaSalle’s theorem [46], customized to the situation that the set 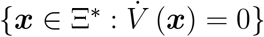 is equal to {***x**_e_*}.

Despite the fact that this manuscript has been devoted to explore systems with continuously differentiable right-hand side functions, we need to recall some concepts and results concerning Filippov’s differential inclusion [47–50], having been particularly developed to analyze nonsmooth systems, since they enable us to study the convergence of solutions using nonsmooth functions. Due to the continuity of ***f*** [48], the system (2) can be replaced with the following Filippov differential inclusion: 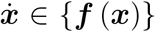. If there is a locally Lipschitz function *V* : ℝ^*M*^ → ℝ that can be written as a pointwise maximum of a set of smooth functions, called a max function, such as ‖***x***‖_1_ = **1**^*T*^ |***x***|, then the derivative of *V* along the trajectories of the differential inclusion 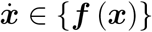 exists almost everywhere and satisfies [49]:

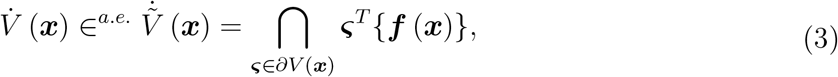

where *∂V* (***x***) represents Clark’s generalized gradient of *V* at point ***x*** which is defined as [51]

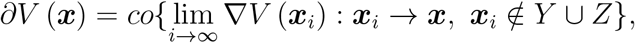

where *co* denotes the convex hull, and *Y* is the set of Lebesgue measure zero where ∇*V* does not exist, and *Z* is also an arbitrary set of zero measure. For example, the function *V* (*x*) = |*x*| with *x* ∈ ℝ has the Clarke generalized gradient

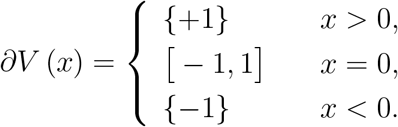

#### Theorem 2.4

(A nonsmooth version of LaSalle’s invariance theorem). *[49] Assume that* Ξ^*\**^ ⊂ *X is a compact set that is positively invariant with respect to Eq. (2). If V* : *X* → ℝ *is a locally Lipschitz max function such that u* ≤ 0 *for all* 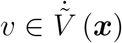 *for all **x*** ∈ Ξ^*\**^ (note that if 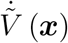 *is the empty set then this condition is trivially satisfied), then every solution in* Ξ*\* converges to the largest invariant set in the closure of 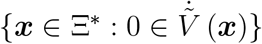*

It is worth noting that the original theorem [49] from which Theorem 2.4 was derived requires uniqueness of solutions, which is automatically established here by Theorem 2.1.

Once a system of differential equations is utilized to describe the temporal evolution of nonnegative variables such as concentrations, the system requires to preserve nonnegativity of solutions for nonnegative initial conditions on the maximal forward time interval of existence of each solution. This property is classically known as positivity, and if a system has it, then it is called a positive system [52]. It is intuitively evident and shown in [53] that the system (2) with 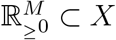 is positive if and only if ***f*** (***x***) satisfies

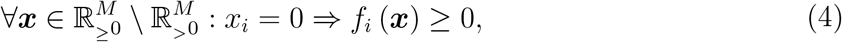

which henceforth will be referred to as the positivity condition.

### 2.3. Weighted directed graphs

A weighted directed graph (or simply a digraph) *G* with no self-loops is a triple (*N_G_, E_G_, ω_G_*), where *N_G_* is a finite nonempty set of nodes (or vertices), *N_G_* = {1, 2, *…, M*} with a positive integer *M*, *E_G_* is a set of directed edges, *E_G_* ⊆ *N_G_* × *N_G_* \ {(*k, i*) ∈ *N_G_* × *N_G_* : *k* = *i*}, and *ω_G_* : *E_G_* → (0, ∞) is a function that associates each edge in *G* from vertex *k* to vertex *i*, *k* → *i*, to a positive real number, *ω_ik_*. If there is no such edge, we let *ω_ik_* = 0. When *E_G_* also satisfies the symmetric condition which implies if (*k, i*) ∈ *E_G_*, then (*i, k*) ∈ *E_G_*, the digraph *G* is here referred to as a symmetric digraph. Note that in our context a weighted undirected graph *G* can be treated as a symmetric digraph when *ω_ik_* = *ω_ki_* for all (*i, k*) ∈ *E_G_*. Given a digraph *G*, a path from node *k* to node *i* is a sequence of successive edges {(*k, k*_1_), (*k*_1_, *k*_2_), *…*, (*k_l_, i*)} ⊆ *E_G_*, i.e.,

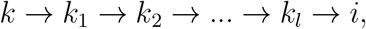

and denoted by 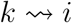, and node *k* is also said to be strongly connected to node *i* if either *k* = *i* or there are both paths 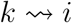 and 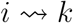. Strong connectivity defines an equivalence relation on the set of nodes, whose equivalence classes are called the strongly connected components (SCCs) of *G*. Now, suppose [*i*]_*SCC*_ and [*k*]_*SCC*_ represent the SCC having the vertex *i* and the SCC containing the vertex *k*, respectively. We say that [*i*]_*SCC*_ precedes [*k*]_*SCC*_, denoted by [*i*]_*SCC*_ ⩽ [*k*]_*SCC*_, if 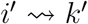 for some *i′* ∈ [*i*]_*SCC*_ and some *k′* ∈ [*k*]_*SCC*_. Since the relation ⩽ on SCCs of *G* is transitive, reflexive, and antisymmetric, it is a partial order, and thus it enables us to determine terminal SCCs of *G*, which are those [*i*]_*SCC*_ such that if [*i*]_*SCC*_ ⩽ [*k*]_*SCC*_ then [*i*]_*SCC*_ = [*k*]_*SCC*_ [54]. Here, 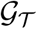 denotes the set of all digraphs whose SCCs are all terminal. It is obvious that symmetric digraphs belong to 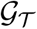.

## 3. Extensions of fractional polynomial functions

Let 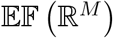 represent the set of all functions from ℝ^*M*^ to ℝ which can be expressed in the form

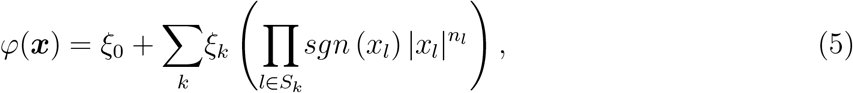

with *ξ*_0_, *ξ_k_* ∈ ℝ, *n_l_* ∈ ℝ_*≥*1_, where the sum is over a finite number of *k*’s, and *S_k_* is a sub-multiset of the multiset {1, 1, 2, 2, *…, M, M*}. A multiset is a set whose elements can be repeated more than once, and the number of times an element appears in a multiset is called its multiplicity [55]. Hence, the multiplicity of each element of the sub-multiset *S_k_* can be at most two. Taking elements with multiplicity 2 into account allows a function 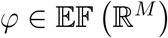 to have terms including 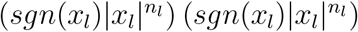, which is equal to 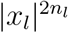, and thus it makes the set 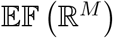 closed under the standard multiplication of functions. Indeed, it can be shown that pointwise addition and multiplication of functions turn 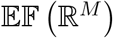 into a commutative ring and also a commutative and associative algebra over ℝ if we define scalar multiplication by (*γφ*) (***x***) ≔ *γφ* (***x***) for any *γ* ∈ ℝ. Note that since proving the mentioned algebraic properties of 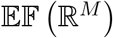 is similar to proving those properties for the set of all continuous functions from ℝ^*M*^ to ℝ [56], we have deliberately avoided discussing their proofs in detail.

The restriction of 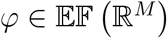 to 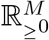 is a FP function. Although FP functions have been used in various fields of research, such as statistical modeling [57, 58], fractional calculus [59, 60], non-integer summations [61–63], and b-function computing [64, 65], to the best of the authors’ knowledge there has been no generalization of a FP function like (5) in previous published studies. The ultimate aim of this manuscript is to develop a framework for describing the spatiotemporal evolution of concentrations of different species, and since the concentration of a given species takes nonnegative values, at first glance it may appear that there would be no reason for proposing the set 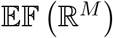, but we have had a theoretical as well as practical motivation behind this generalization. The theoretical reason was to provide continuously differentiable extensions of FP functions on the whole Euclidean space; see Lemma 3.1. Indeed, the conventional definition of a real-valued fractional polynomial function is confined to the nonnegative orthant, which is a closed subset of the Euclidean space, whereas in the literature most theorems and results concerning differentiable functions are only applicable to functions on open domains. To clarify, in any application of functions defined on a closed subset of the Euclidean space, which requires the consideration of their differentiability, it is generally needed either to customize the standard theorems of interest to their closed domains [66] or more typically to find continuous differentiable extensions of these functions to an open subset including their domains [67], due to the fact that closed subsets of the Euclidean space normally lack a differentiable structure. For further investigation about such extensions and their existence, we refer to Whitney’s extension problem [68, 69] and other relevant articles [70–72].

The practical motivation for introducing the set 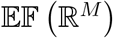 can be explained by defining the *actiue* concentration of a given species that is calculated by subtracting the normalized concentration of its annihilating (or neutralizing) species from its own concentration, where the normalization is done such that zero active concentration corresponds to neutralization. As a result, the active concentration may take negative values, and additionally, if no annihilating species exist, the concept of active concentration is simply equivalent to the usual concept of concentration. In fact, it allows us to simultaneously consider two species that can neutralize the effects of each other at the expense of accepting negative values. Generally speaking, biological studies on species affected by antibodies can benefit from the notion of active concentration. Let us note that although we have introduced the active concentration and will continue our theoretical discussion based on this notion, the simulations in this article are confined to nonnegative initial conditions due to brevity. Indeed, we intend to address more empirical aspects of this work, especially in the context of neurodegenerative diseases, in future articles.

### Lemma 3.1.

*Every function 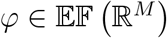 is continuously differentiable on* ℝ^*M*^.

*Proof*. A function *φ* : ℝ^*M*^ → ℝ is said to be continuously differentiable on ℝ^*M*^ if all its partial derivatives exist and are continuous at each point of ℝ^*M*^. Hence, since a function 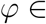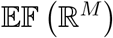 consists of a finite number of additions and multiplications of the terms 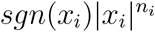, *i* ∈ {1, 2, *…, M*}, and also the constants *ξ*_0_ and *ξ_k_*, it is sufficient, according to the linearity of differentiation and the chain rule, to show that the derivative of 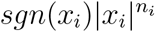 for each *i* ∈ {1, 2, *…, M*} with *n_i_* ∈ ℝ_*≥*1_ exists and is also continuous at every point of ℝ. Using the definition of derivative and applying the L’Hôpital’s rule, it is not difficult to obtain

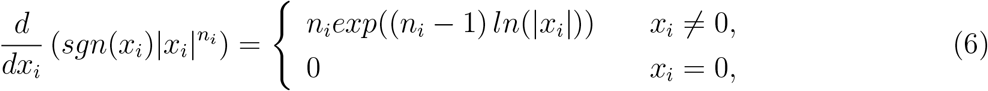

which may be written more compactly as 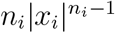. Since it can be easily seen that 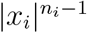 with *n_i_* ≥ 1 is continuous on ℝ, the continuity of the partial derivatives of *φ* on ℝ^*M*^ will be assured.

Let 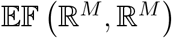 denote the set of all vector-valued functions from ℝ^*M*^ to ℝ^*M*^ whose all *M* real-valued components belong to 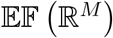. Since a vector-valued function is continuously differentiable on ℝ^*M*^ if and only if all its components are continuously differentiable on ℝ^*M*^, Lemma 3.1 implies the following statement.

### Lemma 3.2.

*Any vector-valued function 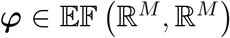 is continuously differentiable on* ℝ^M^.

## 4. Extended FP diffusion equation on directed networks

### 4.1. A generalized nonlinear Laplacian operator on directed networks

We generalize the conventional graph Laplacian operator on a digraph *G* with *M* nodes as follows:

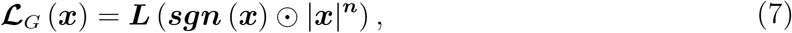

where ***x*** ∈ ℝ^*M*^, and ***n*** is a constant vector belonging to 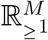, which implies 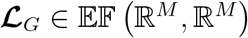. The matrix 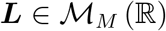 is called the Laplacian matrix and defined by

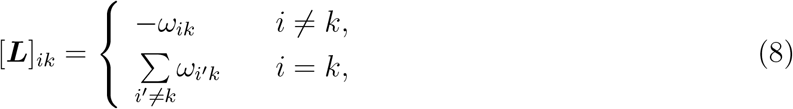

where *ω_ik_* is the weight of the edge *k* → *i*. It can be directly deduced from the definition (8) that the Laplacian matrix satisfies the following equality **1**^*T*^ ***L*** = 0.

#### Lemma 4.1.

*[7, 73] If G is a strongly connected digraph with M nodes, then the kernel of the Laplacian matrix **L**, i.e., ker* (***L***) = {***x*** ∈ ℝ^*M*^ : ***Lx*** = 0}, *is one-dimensional [73], and there is also a vector **u*** ∈ *ker* (***L***) *whose elements are all positive [7]*.

Note that [73] and [7] consider −***L*** as the Laplacian matrix. However, since −***Lu*** = 0 implies ***Lu*** = 0, their results are also applicable to ***L***.

#### Lemma 4.2.

*[7] Assume that 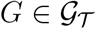 has M nodes and m SCCs. Then, there is a permutation matrix **P** _d_ that transforms the Laplacian matrix **L** of G into a block diagonal form, i.e*.,

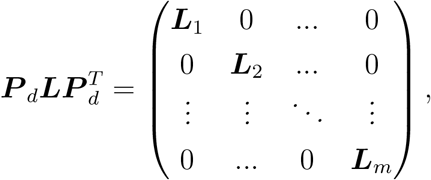

*where every block of 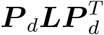 corresponds to a unique SCC of G*.

### 4.2. A Model for Diffusion Processes

Given a digraph 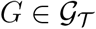 having *M* nodes and *m* SCCs, let us study a diffusion process on *G* that may be described by

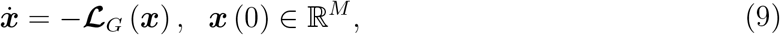

where the *i*-th element of ***x*** (*t*) is the active concentration of a given species at node *i* at time *t*, and accordingly, *x_tot_* (*t*) = **1**^*T*^ ***x*** (*t*) is their total active concentration on *G* at time *t*.

#### Lemma 4.3.

*The total active concentration is preserved during a diffusion process governed by the differential equation (9).*

*Proof*. Using **1**^*T*^ ***L*** = 0, we obtain

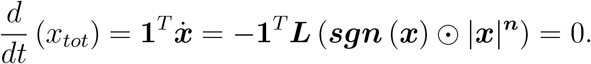

#### 4.2.1. Existence and uniqueness of solutions and positively invariant sets

##### Proposition 4.1.

*The set* ϒ_*η*_ = {***x*** ∈ ℝ^*M*^ : ‖***x***‖ ≤ *η*} *with η* ∈ ℝ_≥0_ *is positively invariant with respect to the system (9)*.

*Proof*. Suppose ***ε*** = (*ε*_1_, *…, ε_M_*)^*T*^ where *ε_i_* is either –1 or 1. Then, we define

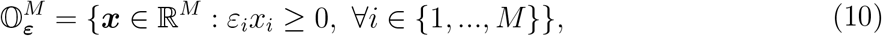

which is obviously an orthant of ℝ^*M*^. Each boundary segment ***ε**^T^ **x*** = *η* of the set ϒ_*η*_ is totally contained in the orthant 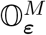, and clearly, it can be seen that ***ε**^T^ **x*** ≥ 0 for 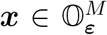 due to *ε_i_x_i_* ≥ 0. Calculating the derivative of ***ε**^T^ **x*** along the trajectories of the system (9) yields

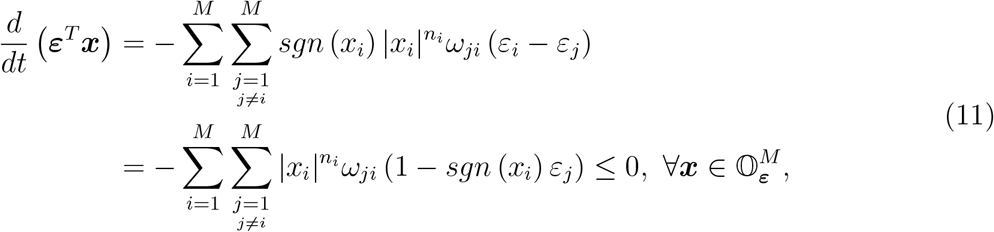

which demonstrates that at any point on each boundary segment ***ε**^T^ **x*** = *η*, ***ε**^T^ **x*** is nonincreasing along the trajectories of the system (9). Therefore, any solution ***x*** (*t*) with an initial condition ***x*** (0) ∈ ϒ_*η*_ cannot leave the set ϒ_*η*_, and the proof is concluded by Theorem 2.1.

##### Corollary 4.1.

*For any initial condition **x*** (0) ∈ ℝ*^M^, the system (9) has a unique bounded solution **x*** (*t*) *that is defined for all t* ≥ 0.

*Proof*. For any initial condition ***x*** (0) ∈ ℝ^*M*^, it is seen that ***x*** (0) ∈ ϒ_*η*_ with *η* ≥ ‖***x*** (0)‖_1_. The rest of the proof follows from Proposition 4.1 and Theorem 2.1.

##### Proposition 4.2.

*The sets*

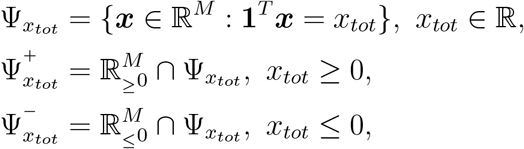

*are positively invariant with respect to the system (9)*.

*Proof*. For the set 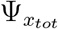, the conclusion follows directly from Lemma 4.3 and Proposition 4.1. Due to the fact that the system (45) is invariant under the change of variables ***x*** → −***x***, its state space is symmetric under reflection through the origin. Hence, it is only sufficient to continue the proof for the set 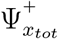. According to Lemma 4.3, any solution ***x*** (*t*) of the system (9) with an initial condition 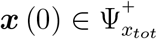 always satisfies **1**^*T*^ ***x*** (*t*) = *x_tot_*. Additionally, ***x*** (*t*) cannot leave the set 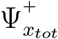 through the boundary of the nonnegative orthant, thanks to the fact that 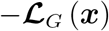 satisfies the positivity condition (4). Thus, the solution ***x*** (*t*) lies entirely in the compact set 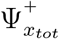 on the maximal forward time interval of its existence. Further, since the right-hand side of Eq. (9) is continuously differentiable on ℝ^*M*^, Theorem 2.1 ensures the existence and uniqueness of the solution ***x*** (*t*) for all *t* ≥ 0. Hence, it can be concluded that

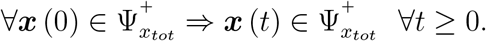

##### Corollary 4.2.

*For any initial state 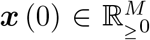 (resp., 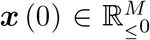), the system (9) has a unique solution **x*** (*t*) *remaining entirely in 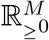 (resp., 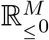) for all t* ≥ 0.

*Proof*. Every solution ***x*** (*t*) of the system (9) with 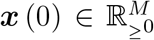 lies entirely in the compact set 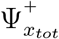 with *x_tot_* = **1**^*T*^ ***x*** (0) by Proposition 4.2. The rest of the proof is similar to that of Proposition 4.2.

Compared to Corollary 4.2, a stronger conclusion can be drawn when *G* is strongly connected.

##### Proposition 4.3.

*Assume G is strongly connected. For any initial condition 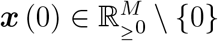 (resp., 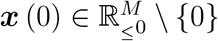), the system (9) has a unique solution **x*** (*t*) *remaining entirely in 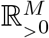 (resp., 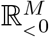) for all t* > 0.

*Proof*. Owing to the fact that the state space is symmetric under reflection through the origin, it suffices only to proceed with the nonnegative orthant. Before we embark on the proof, let us rewrite the system (9) as

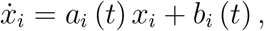

where

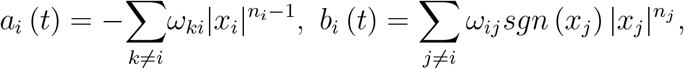

According to the solution formula for a first-order linear differential equation, we obtain

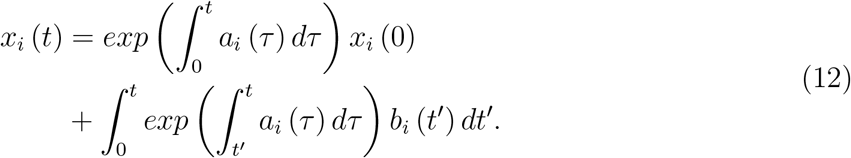

Due to 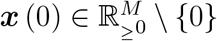, there is at least one component *x_i*_* for which *x_i*_* (0) > 0. Considering Corollary 4.2 and the fact that *ω_ij_* ≥ 0, the initial condition *x_i*_* (0) > 0 implies *x_i*_* (*t*) > 0 for all *t* > 0 using Eq. (12). Since *G* is strongly connected, there is at least one edge from node *i\** to another node *j\**, which leads to *b_j*_* (*t*) > 0 for all *t* > 0. Thus, using again the formula (12), we deduce that *c_j*_* (*t*) > 0 for all *t* > 0. Continuing this reasoning yields 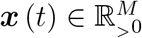 for all *t* > 0.

Recalling Lemma 4.2, it can be demonstrated that the change of variables *y* = ***P** _d_**x*** and ***n**\** = ***P** _d_**n*** transforms the system (9) into

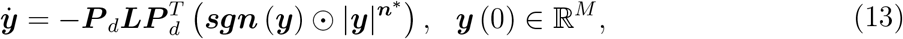

where 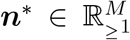 due to the assumption 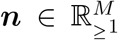. Note that 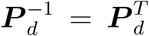, resulting from the fact that any permutation matrix is an orthogonal matrix. It is also easy to see that *x_tot_* = **1**^*T*^ ***x*** = **1**^*T*^ ***y***, and hence the set {***y*** ∈ ℝ^*M*^ : **1**^*T*^ ***y*** = *x_tot_*} is equal to the set 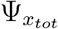, which is positively invariant by Proposition 4.2. Since 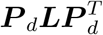 has a block diagonal form, the analysis of a diffusion process on *G* governed by the differential equation (13) can be done for each SCC of *G* independently. Thus, let us rewrite the system (13) as follows:

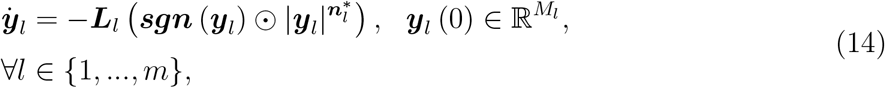

where ***L**_l_* and 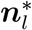 denote a block of 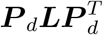 and a subvector of ***n**\** corresponding to the *l*-th SCC of *G*, respectively. *M_l_* is the number of nodes belonging to the SCC *l*, and subsequently, it follows that 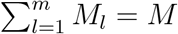 and the Cartesian product 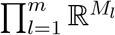 is equal to ℝ^*M*^ by defining the vector 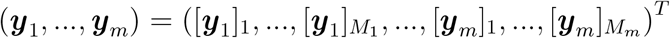 where [***y**_l_*]_*k*_ is the *k*-th element of 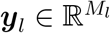. Furthermore, given 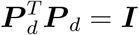, the equality **1**^*T*^ ***L*** = 0 leads to

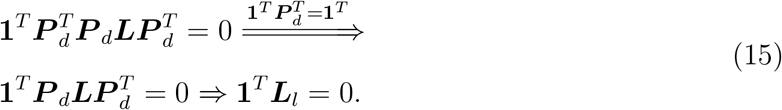

Generally, a fixed value of the total concentration on a digraph belonging to 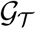 may correspond to more than one equilibrium point of the system (14). In other words, the set 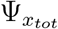 can contain more than one equilibrium point. For example, given a graph *G* with three nodes and with no edges, every point ***y*** ∈ ℝ^3^ is an equilibrium point of the system (14), and as long as the condition 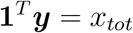 is satisfied, the equilibrium point ***y*** belongs to 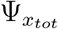. However, in Proposition 4.4 we show that if the total active concentration on each SCC of a digraph 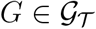 is fixed, then the total active concentration on *G* will correspond to only one equilibrium point of the system (14).

##### Proposition 4.4.

*Consider the system (14) and the sets*

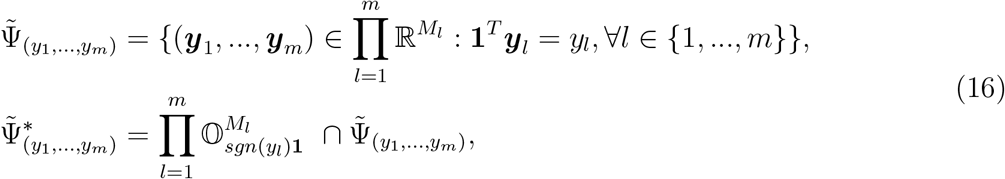

*where the value y_l_ stands for the total active concentration on the l-th SCC of G. Then, each set in (16) contains exactly one equilibrium point of the system (14). Note that 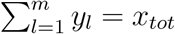 also leads to 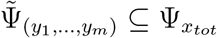.*

*Proof*. Due to brevity, we only provide a proof for the set 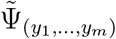. Let ***y**_e_* ∈ ℝ^*M*^ be an equilibrium point of the system (14). Suppose that 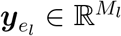 is a subvector of ***y**_e_* corresponding to the SCC *l*. Thus, 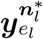 belongs to the kernel of the block ***L**_l_*. Since each component of *G* is strongly connected, Lemma 4.1 implies that *ker* (***L**_l_*) has dimension one, and there is also a subvector ***v**_l_* ∈ *ker* (***L**_l_*) whose elements are positive. Hence, there exists a real nonnegative number *ϱ_l_* such that

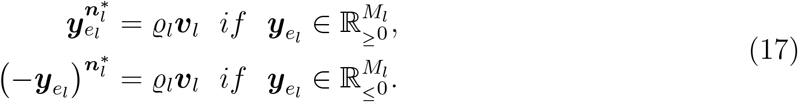

Assume without loss of generality that 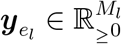, and let us rewrite Eq. (17) as follows:

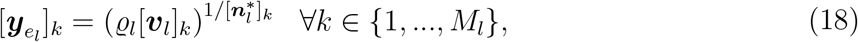

where the index *k* means the *k*-th element. Note that 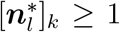 for all *k*. Furthermore, if 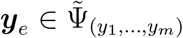, then 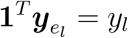, and the substitution of Eq. (18) in 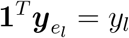 yields

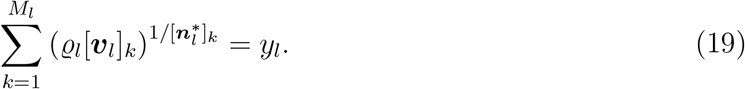

Since the left-hand side of Eq. (19) is a strictly increasing function of *ϱ_l_* for its nonnegative values, there is a one-to-one correspondence between *ϱ_l_* and *y_l_*, and it follows for a fixed value of *y_l_* that there exists only one subvector 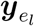 which corresponds to *y_l_*. Therefore, there is a one-to-one correspondence between the vector (*y*_1_, *…, y_m_*)^*T*^ ∈ ℝ^*m*^ and the vector 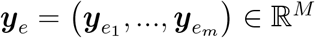, which leads to the conclusion that the set 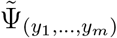 includes exactly one equilibrium point of the system (14).

##### Corollary 4.3.

*Both sets in (16) are positively invariant with respect to the system (14)*.

*Proof*. It can be proven by repeating the proof of the Proposition 4.2 for each SCC of *G*.

#### 4.2.2. Convergence of solutions and stability of equilibria

To investigate the convergence of solutions of the system (14), we first exploit the Lyapunov function candidate

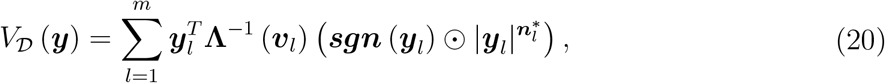

where 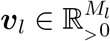 belongs to *ker* (***L**_l_*), and its existence is guaranteed by Lemma 4.1. Note that **Λ**^−1^ (***v**_l_*) denotes the inverse of the matrix **Λ** (***v**_l_*). The function 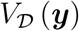 can also be rewritten as follows 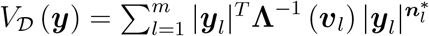, for which it is easy to notice the positive definiteness with respect to ***y*** = 0, i.e.,

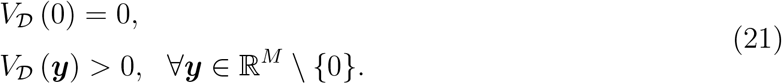

Considering 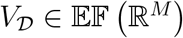, this function is continuously differentiable on ℝ^*M*^ by Lemma 3.1, and the calculation of its derivative along the trajectories of the system (14) yields

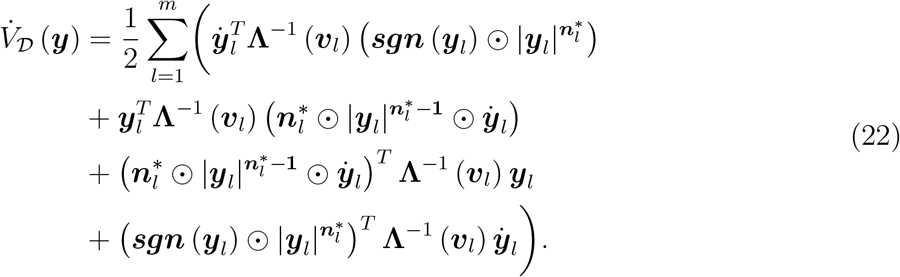

Using the change of variables 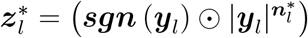, and substituting Eq. (14) into Eq. (22), we have

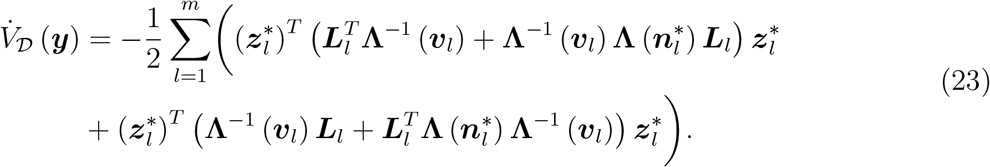

Finally, the change of variables 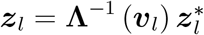 brings Eq. (23) into

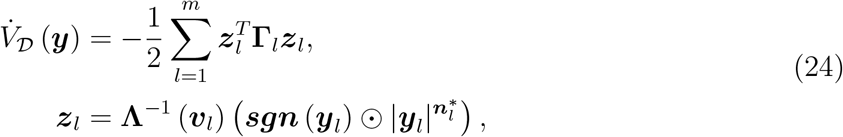

where

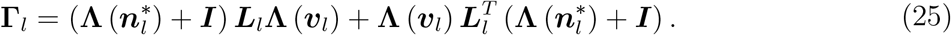

It is easy to spot that **Γ**_*l*_ is a symmetric matrix, and thus to conclude that 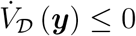 for all ***y*** ∈ ℝ^*M*^ is equivalent to showing that **Γ**_*l*_ for all *l* is a positive semidefinite matrix. If the matrix **Γ**_*l*_ is positive semidefinite, then it must possess the following property: 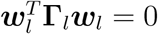 if and only if **Γ**_*l*_***w**_l_* = 0 for all 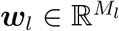. See [44] for a proof of this property. Hence, since the relation ***v**_l_* ∈ *ker* (***L**_l_*) results in **1**^*T*^ **Γ**_*l*_**1** = 0, we must have **Γ**_*l*_**1** = 0, which requires

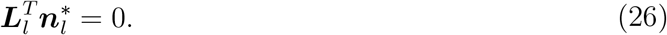

In fact, the equality (26) is a necessary condition for **Γ**_*l*_ to be positive semidefinite. Furthermore, since this equality ensures **Γ**_*l*_**1** = 0, it is also a sufficient condition for **Γ**_*l*_ to be positive semidefinite by Lemma 4.4.

##### Lemma 4.4.

*Let 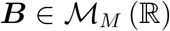 be a symmetric matrix whose off-diagonal elements are nonpositive*.

- *If* 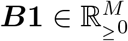, *then **B*** ⪰ 0.
- *If* 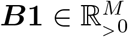, *then **B*** ≻ 0.

*Proof.* It is a corollary of Geršgorin circle theorem. Geršgorin circle theorem states that every eigenvalue *λ* of the matrix ***B*** belongs to the union of Geršgorin discs, i.e.,

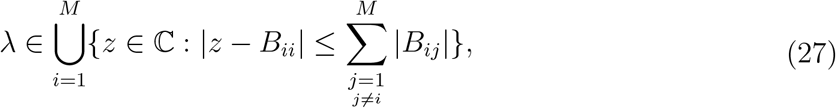

where ℂ and *B_ij_* denote the complex plane and the entry in the *i*-th row and *j*-th column of ***B***, respectively. Since ***B*** is symmetric, all its eigenvalues are real. Thus, given that its off-diagonal elements are also nonpositive, the relation (27) can be rewritten as follows:

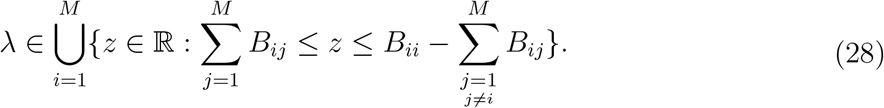

Considering the relation (28), 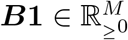 (resp., 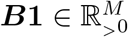) ensures that all eigenvalues of ***B*** are equal to or greater than zero (resp., greater than zero), which confirms that ***B*** is positive semidefinite (resp., positive definite).

Because ***L**_l_* is a square matrix, the dimension of *ker* (***L**_l_*) is equal to the dimension of 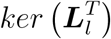, which is a standard result in linear algebra. Hence, given that each component of *G* is strongly connected, Lemma (4.1) implies that 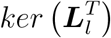 has dimension one. In addition, owing to the fact that 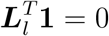 (see Eq. (15)), the condition (26) can boil down to

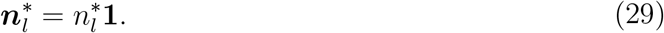

considering 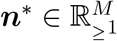, the coefficient 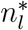 must be greater than or equal to one. An immediate conclusion from the foregoing discussion is that if the system (14) holds the condition (29) for all *l*, then by Theorem 2.2 the function 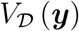 in Eq. (20) is a Lyapunov function to demonstrate the stability of the equilibrium point at the origin, which is of no practical interest in view of the fact that the equilibrium point ***y**_e_* = 0 corresponds to the zero total concentration. However, when the condition (29) is satisfied, Lemma 4.5 reveals more details about the derivative of 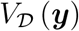 along the trajectories of the system (14) which will assist us in achieving a result concerning the convergence of solutions in the case of an arbitrary total concentration.

##### Proposition 4.5.

*Assume that the system (14) satisfies the condition (29) for all l. Then, for any equilibrium point **y**_e_* ∈ ℝ^*M*^ *and the set* 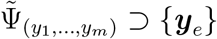, *the function* 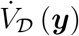 *in Eq. (24) meets the following conditions*:

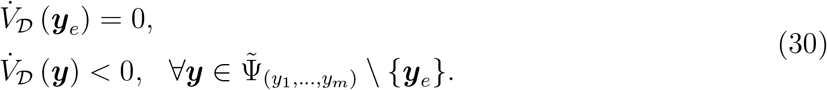

*Proof*. Assume that 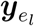 is a subvector of ***y**_e_* corresponding to the *l*-th SCC of *G*. Since ***y**_e_* is an equilibrium point of the system (14), it must satisfy

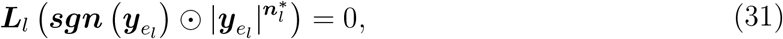

for all *l*, from which it follows that 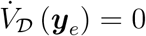. Considering the condition (29), the function 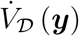 in Eq. (24) can be rewritten as

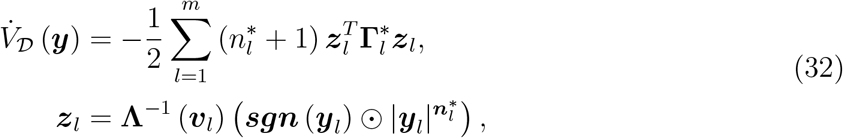

where

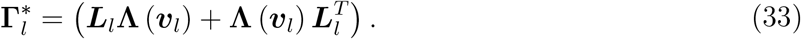

Owing to ***v**_l_* ∈ *ker* (***L**_l_*), we have 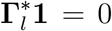, and thus the matrix 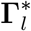 is positive semidefinite by Lemma 4.4. Hence, the function 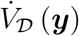 is a summation of nonpositive terms, and it will be sufficient to show that there is at least one nonzero term. Recalling Proposition 4.4, the relation 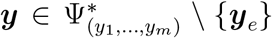 implies that ***y*** is not an equilibrium point of the system (14). Hence, there exists at least one SCC *l′* of *G* such that for the subvector ***y**_l′_* of ***y*** we have 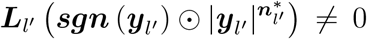 or equivalently ***L**_l′_* **Λ** (***v**_l′_*) ***z**_l′_* ≠ 0; that is to say that ***z**_l′_* ∉ *ker* (***L**_l′_* **Λ** (***v**_l′_*)). In addition, since the matrix 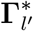 is positive semidefinite, it has the following property: 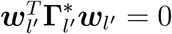 if and only if 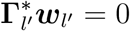 for all 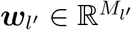. Given ***w**_l′_* = ***z**_l′_*, the conclusion 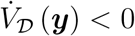 for 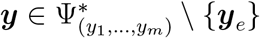 can therefore be drawn from the following equality:

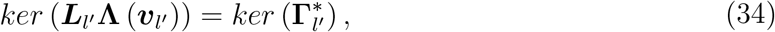

which will be proven as the next step of our proof. The matrix **Λ** (***u**_l′_*) is full rank, deduced from the fact that 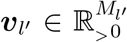. Thus, since *ker* (***L**_l′_*) has dimension one according to Lemma 4.1, it follows that *ker* (***L**_l′_* **Λ** (***u**_l′_*)) is also one-dimensional. Further, it is easily seen that ***u**_l′_* ∈ *ker* (***L**_l′_*) implies **1** ∈ *ker* (***L**_l′_* **Λ** (***u**_l′_*)). Hence, to demonstrate the equality (34), it suffices to show that if 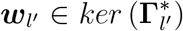, then for elements of ***w**_l′_* we get 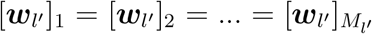. Let 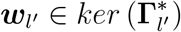, and thus given 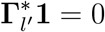, it can be written

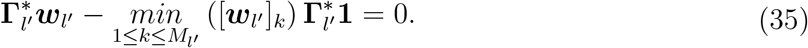

We define a set *U_min_* ⊆ {1, 2, *…, M_l′_*} such that if 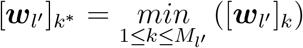, then the index *k\** belongs to *U_min_*. Due to 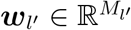, there is at least one element of ***w**_l′_* whose index belongs to *U_min_*, and thus this set is not empty. For *k\** ∈ *U_min_*, we can obtain the following equation using Eq. (35).

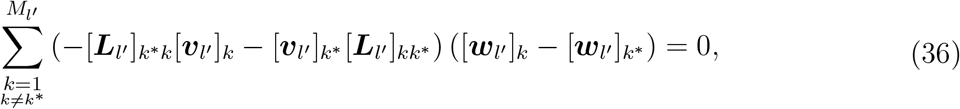

where [***L**_l′_*]_*k*\**k*_ denotes the entry in the *k\**-th row and *k*-th column of ***L**_l′_*. Since 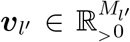,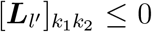 for all *k*_1_ ≠ *k*_2_, and [***w**_l′_*]_*k*_ ≥ [***w**_l′_*]_*k\**_ for all *k*, both terms of the summand in Eq. (36) are nonnegative. Besides, because the component *l′* of *G* is strongly connected, there is at least one edge from *k\** to one of its other nodes. This implies that the first term of the summand in Eq. (36) is in fact strictly positive, and it can be deduced that [***w**_l′_*]_*k*_ = [***w**_l′_*]_*k\**_. As a result, the set *U_min_* is closed under outgoing edges, and hence, we conclude that *U_min_* = {1, 2, *…, M_l′_*}, which means 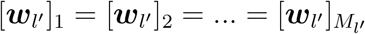.

It is now time to encapsulate our first finding regarding the convergence of solutions of the system (14) in Corollary 4.4.

##### Corollary 4.4.

*Let **y**_e_* ∈ ℝ*^M^ be an equilibrium point of the system (14), holding the condition (29) for all l. Then, for any initial state **y*** (0) *belonging to the positively invariant set 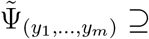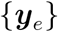 (or 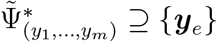), the solution **y*** (*t*) *converges to **y**_e_ as t approaches infinity*.

*Proof*. It is concluded from Theorem 2.3, Corollary 4.3, and Proposition 4.5.

In the case that the system (14) does not fulfill the condition (29), the convergence of solutions to equilibria can also be established in two steps. First, we will demonstrate that every solution of the system (14) with an arbitrary initial condition enters a specific subspace containing equilibrium points. Second, it will be shown that all solutions in that subspace converge to equilibria. The first step is accomplished by using the function *V* (***x***) = ‖***x***‖_1_ whose set-valued derivative, introduced in (3), with respect to the Filippov differential inclusion 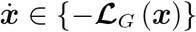 is as follows:

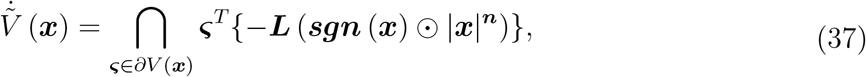

where ***ς*** ∈ *∂V* (***x***) can be expressed elementwise as

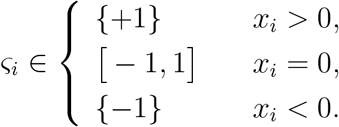

##### Proposition 4.6.

*Consider the differential inclusion 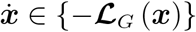 and the set-valued derivative 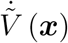 in Eq. (37). If G is strongly connected, then*

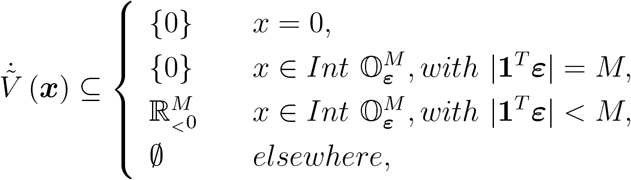

*where 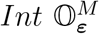 means the interior of the orthant 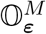, defined in Eq. (10).*

*Proof*. Let us first rewrite Eq. (37) as follows:

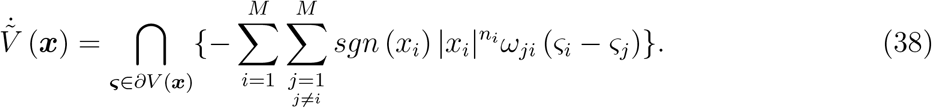

For ***x*** = 0, we have

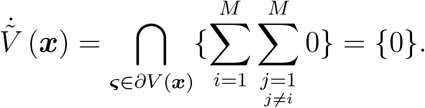

For the case 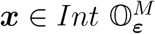 with |**1**^*T*^ **ε**| = *M*, it is obtained that

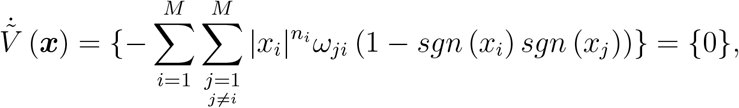

thanks to the fact that *sgn* (*x_i_*) *sgn* (*x_j_*) = 1 for all *i* and *j*.

For the case 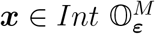 with |**1**^*T*^ ***ε***| < *M*, Eq. (38) can be written as

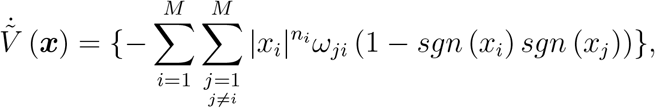

which is obviously nonpositive. Owing to the fact that *G* is strongly connected, it is deduced that 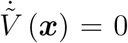 if and only if *sgn* (*x_i_*) *sgn* (*x_j_*) = 1 when *i* ≠ *j*. Indeed, this inference comes from the fact that there is a path passing through all nodes of *G*. Furthermore, since *G* is strongly connected, it follows from |**1**^*T*^ **ε**| < *M* that there is an edge *i\** → *j\** for which *sgn* (*x_i*_*) *sgn* (*x_j*_*) = −1. Thus, we have 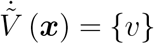 with *u* < 0.

In the last case, ***x*** is a nonzero vector possessing at least one zero element. Hence, due to the fact that *G* is strongly connected, there is an edge *i\** → *j\** for which *x_i*_* ≠ 0 and *x_j*_* ≠ 0, and subsequently, it follows from Eq. (38) that

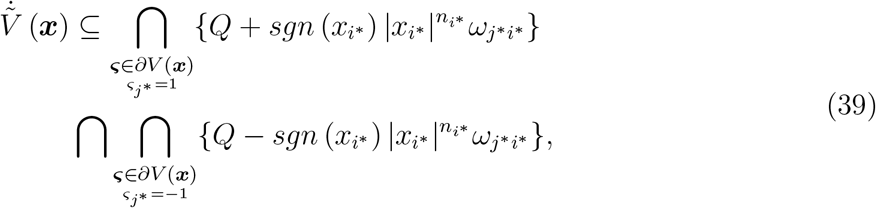

where

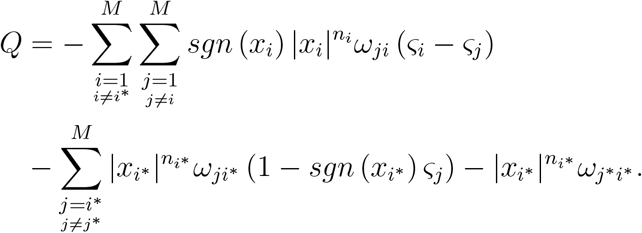

According to (39), it is inferred that 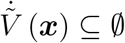 since *x_i_* ≠ 0 and *ω_j*i*_* > 0.

##### Corollary 4.5.

*If G is strongly connected, then every solution of the system (9) with an arbitrary initial state converges to the largest invariant set in the union of 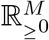 and 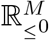.*

*Proof*. It follows from Propositions 4.1 and 4.6 and Theorem 2.4 that a solution ***x*** (*t*) of the system (9) with an initial state ***x*** (0) ∈ ℝ^*M*^ converges to the largest invariant set in the union of 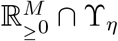 and 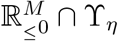 where *η* ≥ ‖***x*** (0)‖_1_.

To prove that all solutions starting in 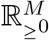 or 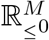 converge to equilibria, we will utilize a modified version of the “free-energy” function of the Becker-Döring model, describing the evolution of coagulation and fragmentation of clusters [74, 75]. Buhagiar [74] observed that this free-energy function is a Lyapunov function for the Becker-Döring cluster equations, and recently, it has been demonstrated that it is also a Lyapunov function for any system of differential equations generated by chemical reaction networks whose components are strongly connected [32]. Due to the fact that it is possible to derive a system of differential equations similar to Eq. (9), where all elements of the power vector ***n*** are nonnegative integers, from a chemical reaction network in which each reaction has identical reactants and also yields identical products, the mentioned free-energy function has assisted us to suggest the function 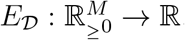, defined by

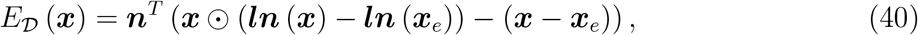

to assess stability properties of an equilibrium point 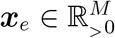 and the convergence of solutions of the system (9). It follows from Lemma 4.5 that 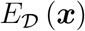 is continuous on 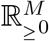 as well as positive definite with respect to ***x**_e_*.

##### Lemma 4.5.

*Assume that E* (*x*) = *x* (*ln* (*x*) – *ln* (*y*)) – (*x* – *y*) *with x* ∈ ℝ_*≥*_0 *and y* ∈ ℝ_>_0 *where E* (0) *is defined to be y. Then*,

- *E* (*x*) *is continuous on* ℝ_*≥*_0;
- *E* (*x*) *is continuously differentiable on* ℝ_>_0;
- *E* (*x*) > 0 *for all x ≠ y, and E* (*y*) = 0.

*Proof*. Considering lim_*x→*0+_ *E* (*x*) = *y*, the first item is easy to spot. Due to *dE/dx* = *ln* (*x*) – *ln* (*y*), the second item is also obvious. Since *d*^2^*E/dx*^2^ = 1/*x* > 0 for all *x* > 0, the function *E* (*x*) is strictly convex on ℝ_>_0, and thus, for all *x, y* ∈ ℝ_>_0 when *x* ≠ *y*, we have

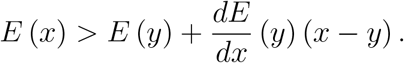

Owing to the fact that *E* (*y*) = (*dE/dx*) (*y*) = 0 and *E* (0) = *y* > 0, the third item is deduced.

Calculating the derivative of 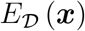 along the trajectories of the system (9) yields

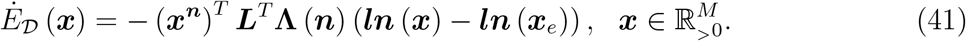

Considering the equalities

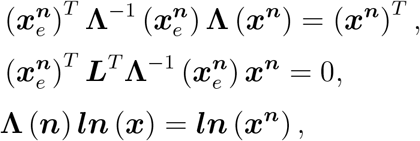

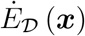 in Eq. (41) can be written as

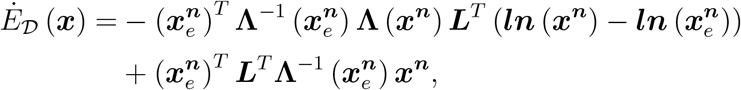

or as the elementwise summation

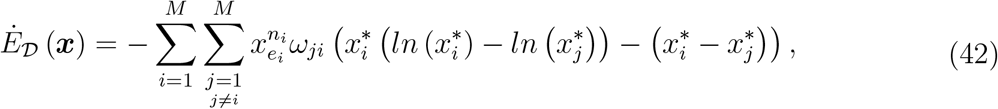

where 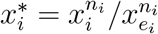

##### Lemma 4.6.

*If G is strongly connected, then*

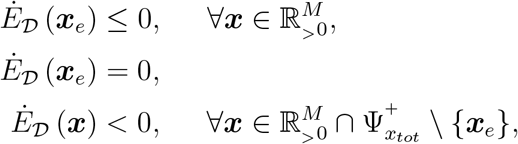

*where x_tot_* = **1**^*T*^ ***x**_e_*.

*Proof*. Due to 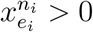 and *ω_ji_* ≥ 0, Lemma 4.5 implies that 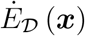 in Eq. (42) is nonpositive for 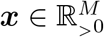. In addition, since *G* is strongly connected, the equality 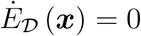 occurs if and only if

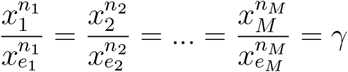

or equivalently 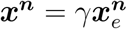 where *γ* > 0 using Lemma 4.5. If ***x*** simultaneously satisfies 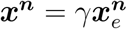 and **1**^*T*^ ***x*** = **1**^*T*^ ***x**_e_*, then we must have

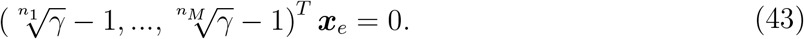

Due to *n_i_* ≥ 1 and 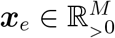, the equality (43) is only satisfied for *γ* = 1.

##### Proposition 4.7.

*Let 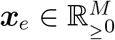 (resp., 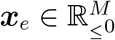) be an equilibrium point of the system (9). If G is strongly connected, then **x**_e_ is stable, and every solution **x*** (*t*) *with an initial state belonging to 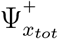 (resp., 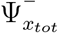) with x_tot_* = **1**^*T*^ ***x**_e_ converges to **x**_e_ as t approaches infinity*.

*Proof*. It is sufficient to proceed only with 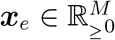 since the state space is symmetric under reflection through the origin. When *G* is strongly connected, we have either ***x**_e_* = 0 or 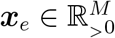 by Lemma 4.1. For the trivial case ***x**_e_* = 0, the stability can be guaranteed by Proposition 4.1. Now, suppose that 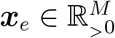. It follows from Lemmas 4.5 and 4.6 that there exists a domain 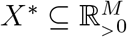 containing ***x**_e_* in which 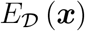 and 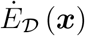 satisfy the conditions of Theorem 2.2. Hence, ***x**_e_* is stable. According to Proposition 4.2, the compact set 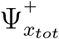 is positively invariant. Now, we show that the function 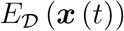 on ℝ_*≥*_0 is decreasing for all 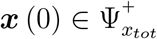 with *x_tot_* = **1**^*T*^ ***x**_e_*. Note that ***x*** (0) ≠ 0 because of *x_tot_* > 0. Due to the fact that the right-hand side of the system (9) is continuously differentiable, any initial condition 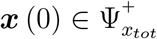 leads to a continuously differentiable solution 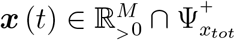 for all *t* > 0 using Proposition 4.3, and since the function 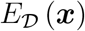 and its derivative 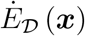 are continuous on 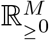 and 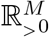, respectively, we can apply the second fundamental theorem of calculus (FTC) as follows:

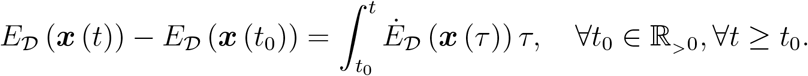

Because 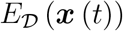 is continuous on ℝ_*≥*_0, we obtain

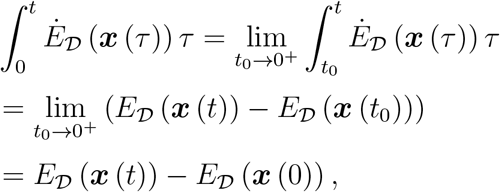

and using Lemma 4.6, which implies 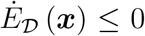 for all 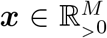, it can be deduced that for every initial state 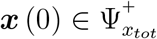, we have

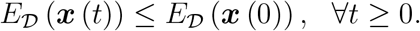

Therefore, 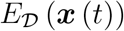 is decreasing, and since 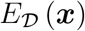 is also lower bounded on the compact set 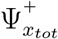 due to its continuity, the function 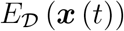 converges to a limit as time goes to infinity, i.e.,

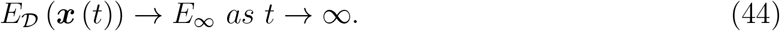

Before going on, we want to recall two definitions. A point ***p*** is said to be in the positive limit set of a solution ***x*** (*t*) if there is a sequence *t_n_* going to infinity with *n* and such that ***x*** (*t_n_*) → ***p*** as *n* → ∞. A set is called invariant if for any initial condition ***x*** (0) belonging to this set, the solution ***x*** (*t*) remains in it for all *t* ∈ ℝ. Now, let us resume our discussion. At this point, someone thinks of applying LaSalle’s invariance principle [46] to investigate the convergence problem here. Owing to the fact that 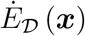 is only continuous on 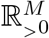 but not 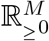, the function 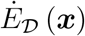 is not continuous on whole 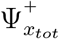, and consequently, LaSalle’s invariance theorem cannot be directly employed. However, we can take advantage of a fundamental result concerning positive limit sets which states that if a solution ***x*** (*t*) is bounded for all *t* ≥ 0, then its positive limit sets is nonempty, compact, and invariant [46]. Let *H*^+^ be the positive limit set of a solution ***x*** (*t*) with 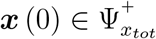. Since the positively invariant set 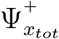 is closed, it follows that 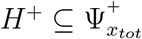. According to (44), it is deduced that

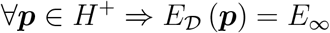

because 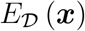 is continuous on 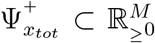. Consider a solution ***x**\** (*t*) with ***x**\** (0) ∈ *H*^+^. Thanks to the fact that *H*^+^ is invariant, it is inferred that 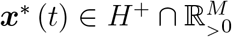 for all *t* > 0 by Proposition 4.3. Thus, recalling again the second FTC results in

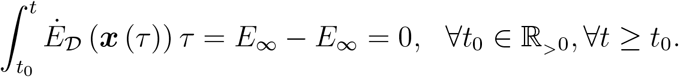

Hence, since 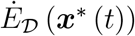 is continuous on [*t*_0_, *t*_1_] for all *t*_1_ > *t*_0_, applying the first FTC implies

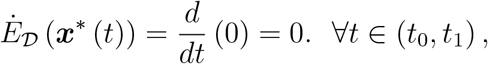

and it follows that ***x**\** (*t*) = ***x**_e_* because of the fact that 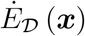 is only zero at ***x**_e_* and is negative for all 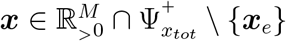 by Lemma 4.6.

We now summarize our findings concerning the convergence of solutions of the system (14) as well as the stability of its equilibria in Corollary 4.6.

##### Corollary 4.6.

*Suppose **y**_e_* ∈ ℝ^*M*^ *is an equilibrium point of the system (14). Then, **y**_e_ is stable, and for any initial state **y*** (0) *belonging to the positively invariant set 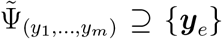 (or 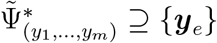), the solution **y*** (*t*) *converges to **y**_e_ as t approaches infinity*.

*Proof*. Considering Proposition 4.4, this conclusion can be drawn by applying Corollary 4.5 and Proposition 4.7 to each subsystem of the system (14).

## 5. Extended FP Fisher-KPP reaction–diffusion equation on directed networks

Given a digraph 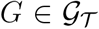 having *M* nodes, let us study a reaction-diffusion process on *G* that may be described by

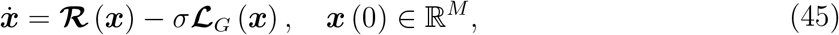

with

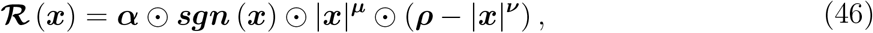

where 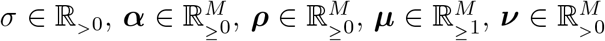, which implies 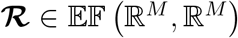, and the *i*-th element of ***x*** (*t*) is the concentration of a given species at node *i* at time *t*, denoted by *x_i_* (*t*). Accordingly, *x_tot_* (*t*) = **1**^*T*^ ***x*** (*t*) is the total concentration of that species on *G* at time *t*.

### Lemma 5.1.

*Let 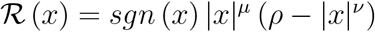 with ρ* ∈ ℝ_*≥*_0, *μ* ∈ ℝ_*≥*_1, *and ν* ∈ ℝ_>_0. *Then*,

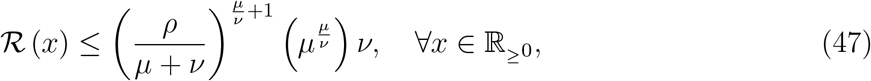

*and*

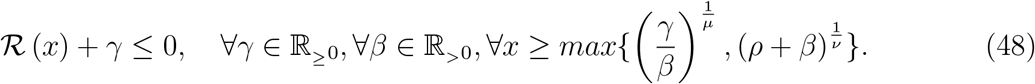

*Proof*. The inequality (47) can be verified by the first derivative test. The derivative of 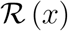 with respect to *x* is given by

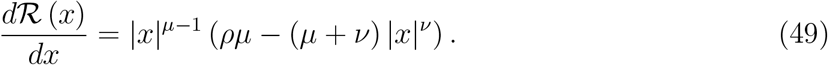

For *ρ* > 0, it is easy to spot that 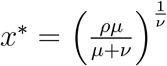 is a critical point of 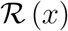, and since we have 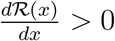 for all *x* between zero and *x\** and 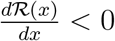 for all *x* greater than *x\**, it follows that 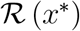, which is equal to the right-hand side of the inequality (47), is the maximum value of 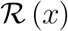 in ℝ_*≥*_0. For the case *ρ* = 0, it is obvious that 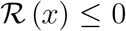 for *x* ≥ 0. The inequality (48) can also be deduced by the following steps:

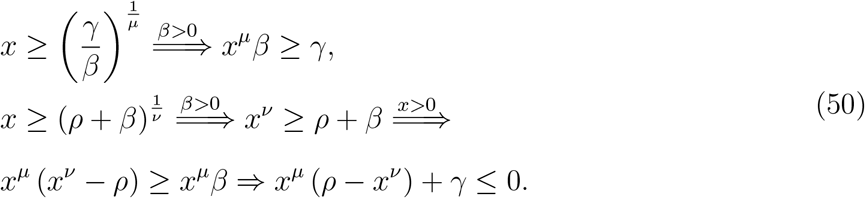

### Proposition 5.1.

*For the system (45), let us define the sets*

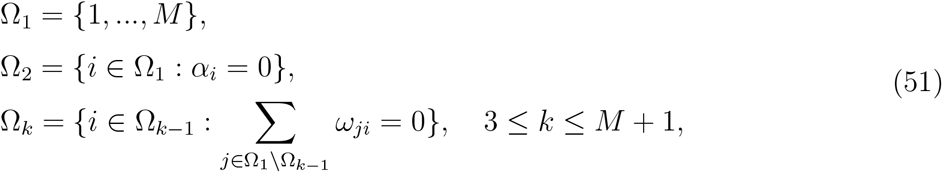

*where α_i_ is the i-th element of **α**, and let M* be the smallest positive integer for which* Ω_*M*_*+1 = Ω_*M*_* *or* Ω_*M*\*+1_ = ∅. *Then, the sets*

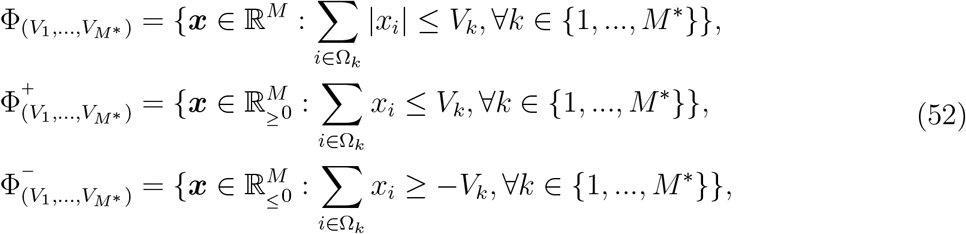

*with 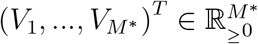 is positively invariant with respect to the system (45) if V_k_’s satisfy for M\** = 1 *that*

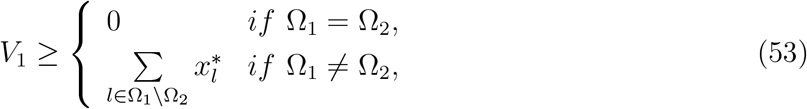

*and for M\** ≥ 2 *that*

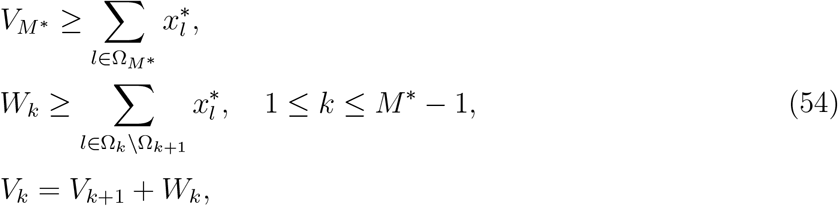

*where, for l* ∈ Ω_1_ \ Ω_2_,

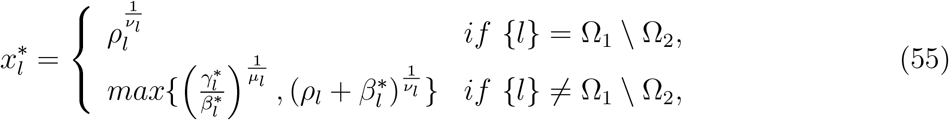

*with*

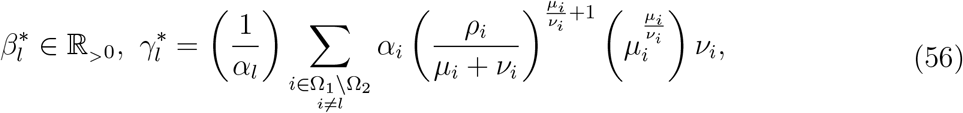

*and, for l* ∈ Ω_*k*_ \ Ω_*k*+1_ *with k* ≥ 2,

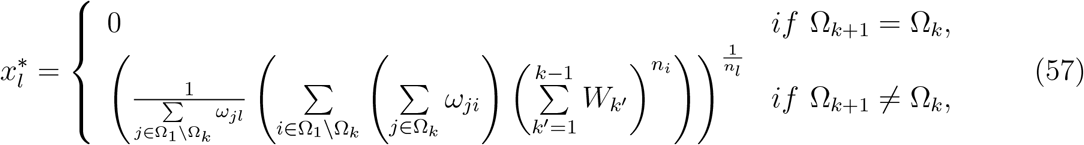

*where ρ_i_, μ_i_, ν_i_, and n_i_ represent the i-th elements of **ρ**, **μ**, **ν**, and **n**.*

*Proof*. Here, we will provide a proof for the set 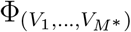, and Consequently, since the right-hand side of Eq. (45) satisfies the positivity condition (4), it can be inferred that 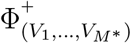 is also positively invariant. Further, thanks to the fact that the system (45) is invariant under the change of variables ***x*** → –***x***, we arrive at the same conclusion for 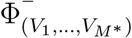.

Assume that ***ε*** = (*ε*_1_, *…, ε_M_*)^*T*^ where *ε_i_* is either −1 or 1. Each boundary segment 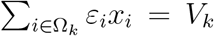 of the set 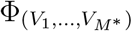 is entirely contained in the orthant 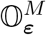, defined in Eq. (10), and it is easy to see that 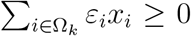 for 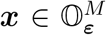 due to *ε_i_x_i_* ≥ 0. Thus, it is sufficient by Theorem 2.1 to demonstrate that at any point 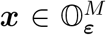 on the boundary segment 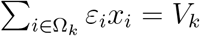, the summation 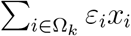 is nonincreasing along the trajectories of the system (45); that is,

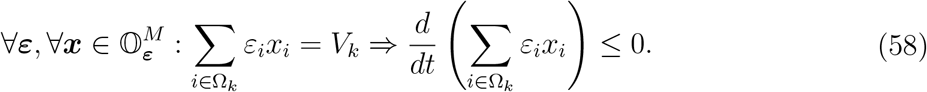

In other words, we intend to show that any solution ***x*** (*t*) with an initial state 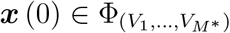 cannot leave the compact set 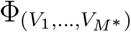 through the boundary segments 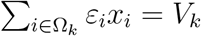, for all *k* ∈ {1, *…, M\**}.

Before we establish the claim (58), we verify the statements (59) and (60) which will be used in the proof. For all *k* ∈ {1, *…, M\**} and *k′ > k*,

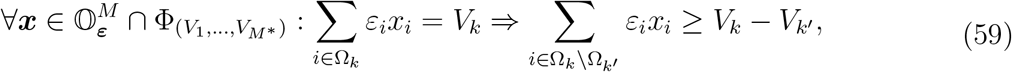

and

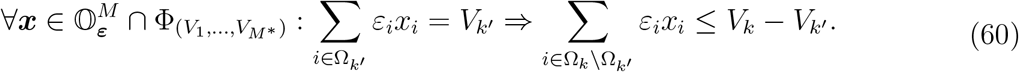

Note that Ω_*k′*_ ⊆ Ω_*k*_ if *k′ > k* by definition. The statement (59) can be proved by contradiction, for if it was not true there would exist 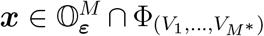 such that 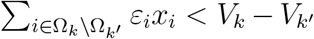. Using 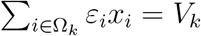, we obtain

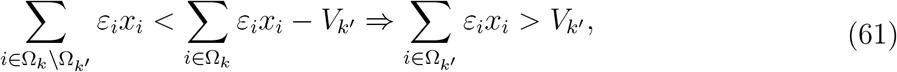

which contradicts the fact that 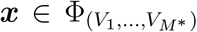. The statement (60) can also be shown by contradiction. Suppose that there is a point 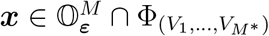 such that 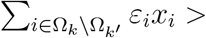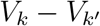. Using 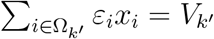, we have

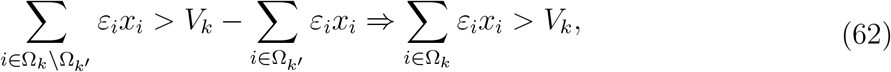

which contradicts the fact that 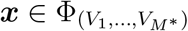.

To prove the claim (58), we start by calculating the derivative of 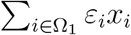 along the trajectories of the system (45).

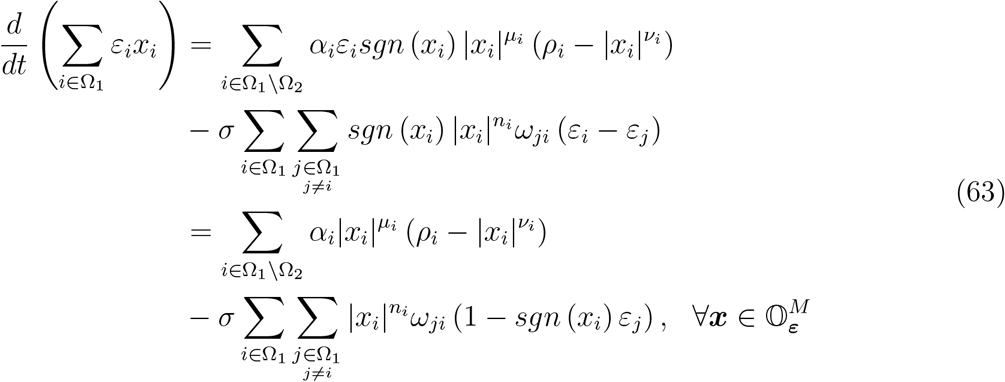

In the case Ω_1_ = Ω_2_, which implies *M\** = 1, the derivative 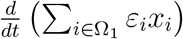 in Eq. (63) is nonpositive for all 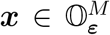 satisfying 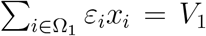 with *V*_1_ ≥ 0; see Eq. (11). In fact, Ω_1_ = Ω_2_ means all reaction coefficients *α_i_*’s are equal to zero, which makes the system (45) equivalent to (9).

In the case Ω_1_ ≠ Ω_2_ when *M\** = 1, we have Ω_2_ = ∅ by definition, and it can be seen that the derivative 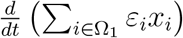 is nonpositive for all 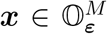 satisfying 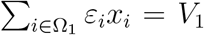 with 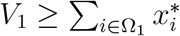. To clarify, when Ω_1_ has only one element *x*_1_, it is easy to see that the derivative 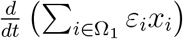 is nonpositive for 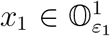 if 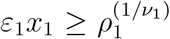, and when Ω_1_ has more than one element, we obtain, for 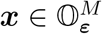 and *l* ∈ Ω_1_ \ Ω_2_,

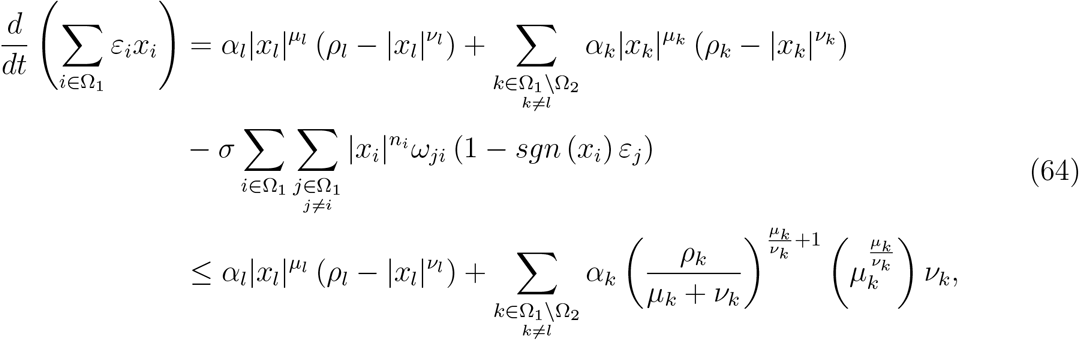

using Lemma 5.1 and Eqs. (63) and (11). Hence, Lemma 5.1 guarantees that the derivative 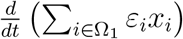 in Eq. (64) is nonpositive for all 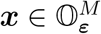 if

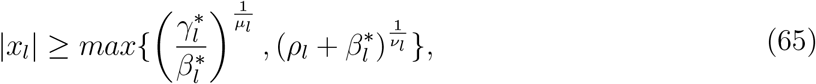

where 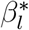 and 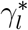 have been given by Eq. (56). Finally, due to 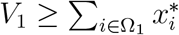, there is at least one component *x_l_* of 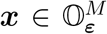 satisfying 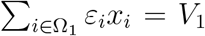 such that 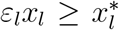, which can be shown by contradiction. Suppose 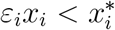 for all *i* ∈ Ω_1_, then 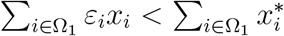, which contradicts the assumption that 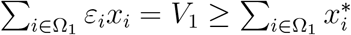.

In the case *M\** ≥ 2, evaluating that the derivative 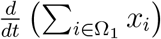 in Eq. (64) is nonpositive for all 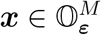 satisfying 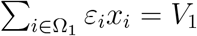 is equivalent by the statement (59) to evaluating that it is nonpositive for all 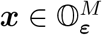 satisfying 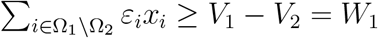, which can be concluded if 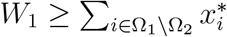. Due to 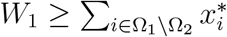, there exists at least one component *x_l_* of 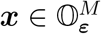 satisfying 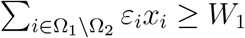 such that 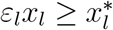, which can be shown by contradiction similar to the previous argument.

To study the other boundary segments 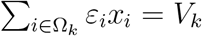 with *k* ≥ 2, we first calculate the derivative of 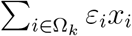, with *k* ≥ 2, along the trajectories of the system (45).

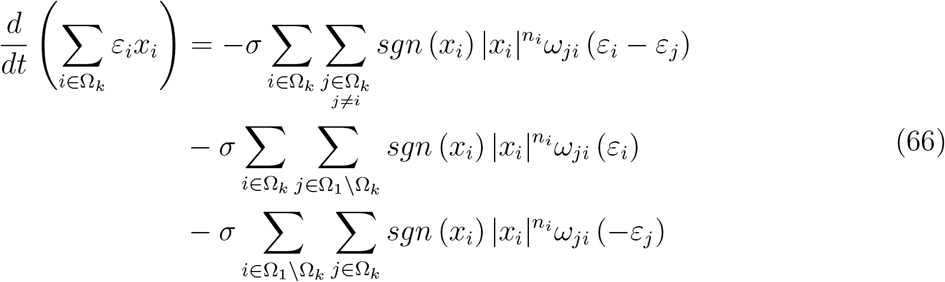

Considering Ω_*k*+1_ ⊆ Ω_*k*_, Eq. (66) can be rewritten as

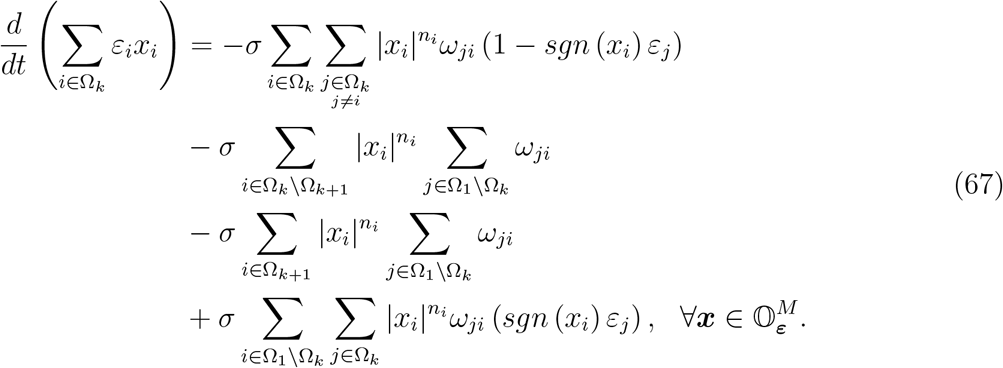

By the definition of Ω_*k*_, the third summation in the right-hand side of Eq. (67) is identical to zero, and thus, we can obtain

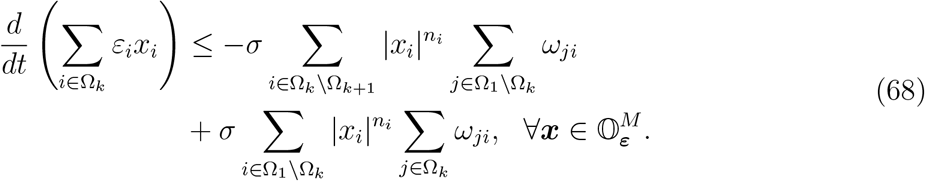

When Ω_*k*+1_ = Ω_*k*_, it can be demonstrated that 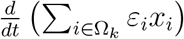 is nonpositive for all 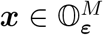. Assuming Ω_*k*+1_ = Ω_*k*_, we have

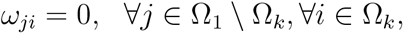

due to *ω_ji_* ≥ 0 and consequently, since all SCCs of *G* are terminal, it is deduced that

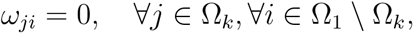

which leads to the conclusion that 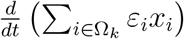 is nonpositive for 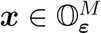 if Ω_*k*+1_ = Ω_*k*_, using the inequality (68). Now supposing that Ω_*k*+1_ ≠ Ω_*k*_, it can be seen for the inequality (68) and *l* ∈ Ω_*k*_ \ Ω_*k*+1_ that

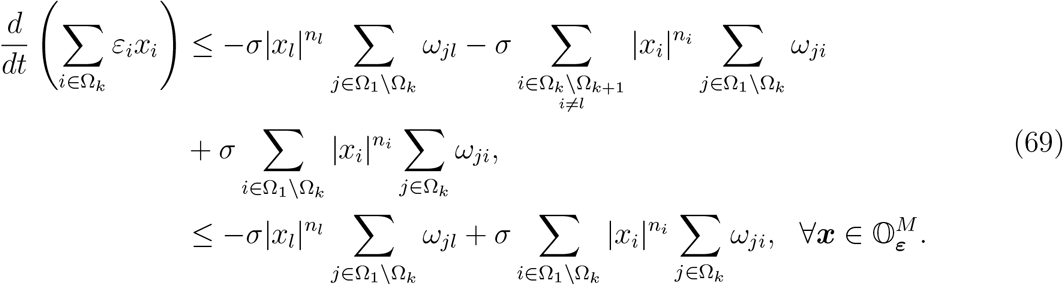

Verifying that the derivative 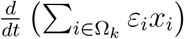 for 2 ≤ *k* ≤ *M\**–1 (resp., *k* = *M\**) is nonpositive for all 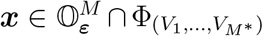 satisfying 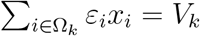 is equivalent by the statements (59) and (60) (resp., the statement (60)) to verifying that it is nonpositive for all 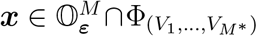 satisfying both 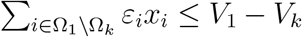 and 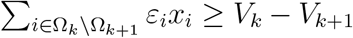 (resp., 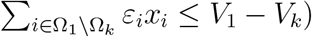. Thus, due to 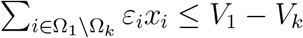, it follows from Eq. (69) that

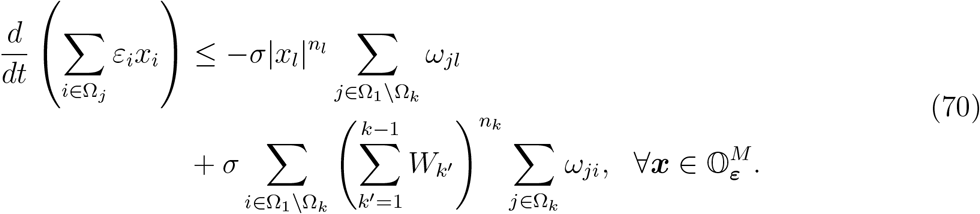

where *W_k′_* = *V_k′_* – *V_k′_*+1. Note by definition that 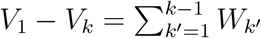. Hence, it is easy to see that 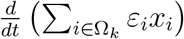 in the inequality (70) is nonpositive if 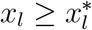 where 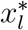 is given in Eq. (57) when Ω_*k*+1_ ≠ Ω_*k*_. Using a contradiction argument similar to the previous one, it can be shown that 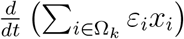 with 2 ≤ *k* ≤ *M\** –1 (resp., *k* = *M\**) is nonpositive for 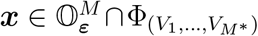 satisfying 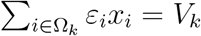 if 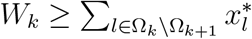 (resp., 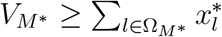).

### Corollary 5.1.

*For any initial condition 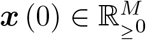 (resp., 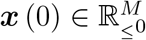), the system (45) has a unique solution **x*** (*t*) *remaining entirely in 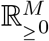 (resp., 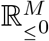) for all t* ≥ 0.

*Proof*. Since the state space is symmetric under reflection through the origin, we only proceed with the nonnegative orthant. For every solution ***x*** (*t*) of the system (45) with 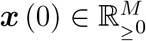, we can choose a vector (*V*_1_, *…, V_M*_*)^*T*^ such that a set 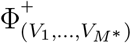 satisfying the conditions of Proposition 5.1 and also 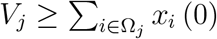 for all *j* ∈ {1, *…, M\**}, which implies 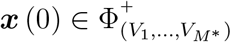. Hence, Theorem 2.1 ensures the existence and uniqueness of the solution ***x*** (*t*) for all *t* ≥ 0.

### Corollary 5.2.

*For any initial condition **x*** (0) ∈ ℝ^*M*^, *the system (45) has a unique bounded solution **x*** (*t*) *that is defined for all t* ≥ 0.

*Proof*. According to the definition (52) of the set 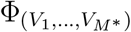, we can find a vector 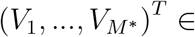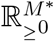 such that 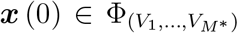 by choosing sufficiently large *V_k_*’s. The rest of the proof follows from Proposition 5.1 and Theorem 2.1.

## 6. Nonlinear diffusion on a directed one-dimensional lattice

Cytoskeletal motor proteins are responsible for directional transportation along microfilaments or microtubules within the cell. Among the prominent motor proteins associated with microtubules, kinesins crawl anterogradely, whereas dyneins slide retrogradely along microtubules by converting the chemical energy produced from ATP hydrolysis to mechanical energy [76]. It was experimentally demonstrated that the movement of microtubule-associated motors can be an anomalous subdiffusion process, especially at long times [77]. In addition, to investigate the motion of motor proteins, recent studies have modeled microtubules as one-dimensional lattices [78–80]. Hence, it may be of potential interest to examine nonlinear diffusion processes governed by the proposed equation (9) on a directed one-dimensional lattice.

Consider a directed one-dimensional lattice *G* with *M* nodes and weights

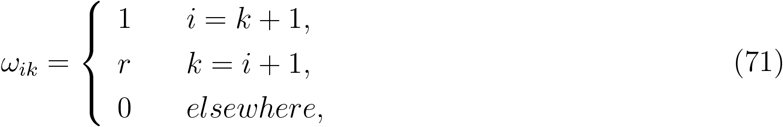

where 0 < *r* ≤ 1. Here, we conduct a qualitative study of particle diffusion on *G* governed by

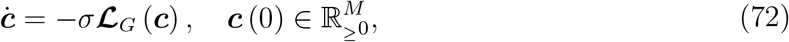

where *σ* ∈ ℝ_>_0 and ***n*** = *n***1** for the Laplacian operator given in Eq. (7). Fig. 1 displays the simulation results of Eq. (72) with *M* = 101 at 10 different points in the parameter space when the initial concentration at each node was set to zero except either for Node 14 or for Node 51, which has been set to 1. As demonstrated by the first and last columns of images in Fig. 1, decreasing *r* results in an anterograde-bias in the diffusion pattern. Furthermore, as seen in the second and third columns of images in Fig. 1, while both shrinking *σ* and growing *n* contribute to more localized diffusion over the simulation period, the tail of the distribution of particles diffused under a regime with an increased *n* is sharper and smaller compared to that of particles diffused under a regime with a reduced *σ*.

**Figure 1:**
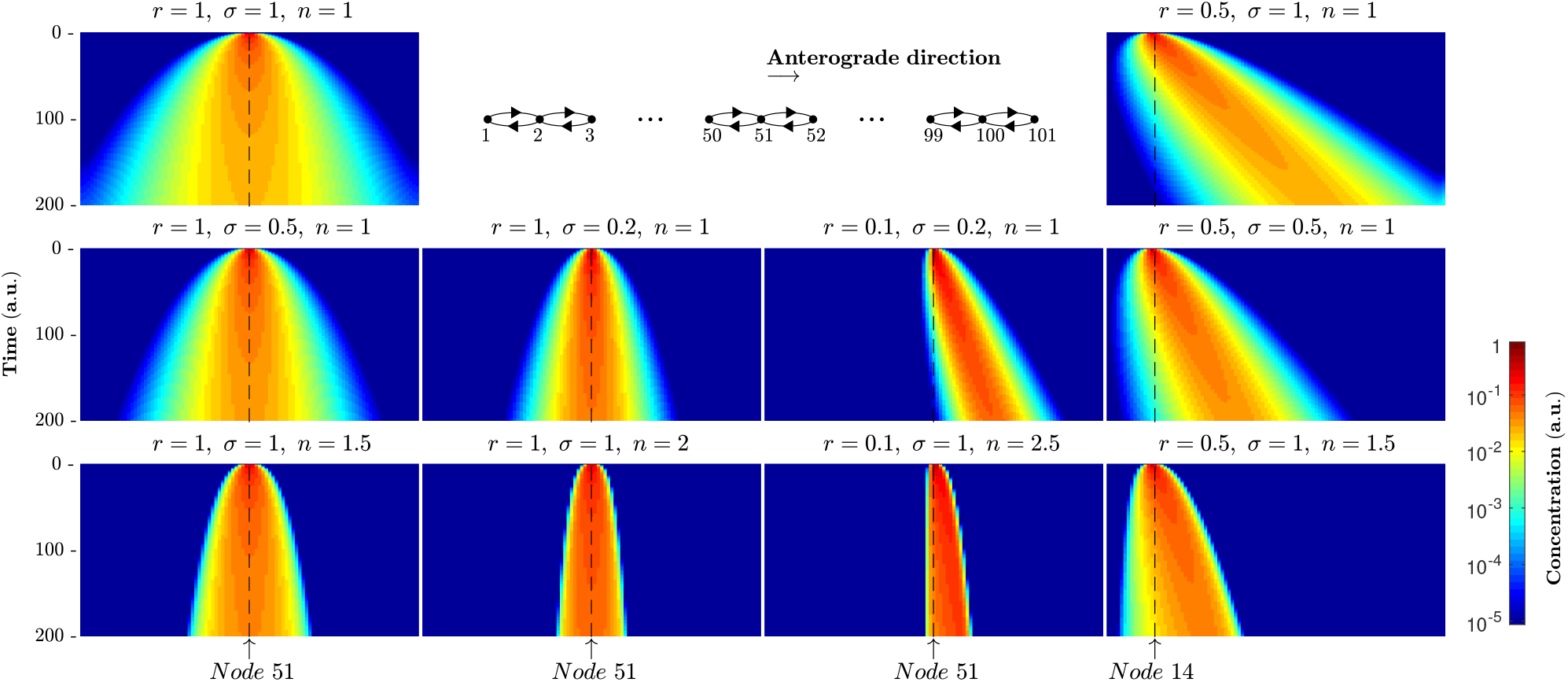
It illustrates a directed one-dimensional lattice with 101 nodes and the simulation results of Eq. (72) for 10 different scenarios when the initial concentration at each node was set to zero except either for Node 14 or for Node 51, which has been set to 1.

Using simulation, we will now demonstrate that, regardless of directionality, diffusion is normal if *n* = 1, and subdiffusion occurs when *n* > 1, but let us first recall two definitions pertaining to anomalous diffusion. The average position of *N* particles and the variance of the position or mean squared displacement with respect to the average position are as follows [81]:

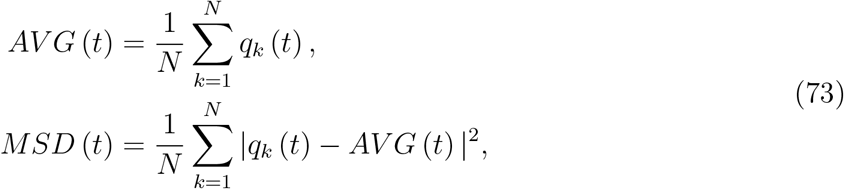

respectively, where *q_k_* (*t*) is the position of the *k*-th particle at time *t*. Assuming *c_i_* denotes the concentration of particles at node *i*, the probability of finding particles at node *i* can be represented as *c_i_*/**1**^*T*^ ***c***, and subsequently, some calculations leads to the formulae

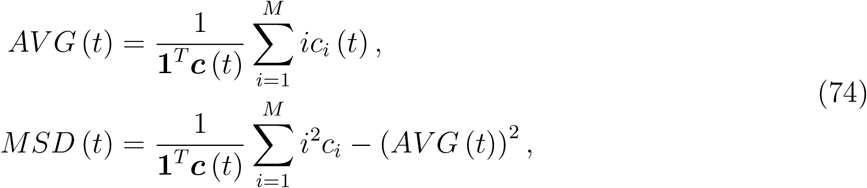

which are simple the mean and the variance of a discrete stochastic process on the set of *M* nodes. Simulation results for two directionally unbiased regimes (*r* = 1) and two anterograde-biased regimes with *r* = 0.1 when the initial concentration at all 101 nodes was set to zero except node 51, which was set to 1, are depicted in Fig. 2. For both directionally unbiased and anterograde-biased regimes with *n* = 1, there is a linear relationship between *MSD* (*t*) and time *t*, indicating normal diffusion, and for the other two regimes with *n* > 1, *MSD* (*t*) is approximately proportional to *t^ζ^* with *ζ* < 1, exhibiting subdiffusion. See Fig. 2A2,B2. Note, however, that this conclusion is confined to a finite time interval before particles accumulate on the lattice’s end points. Moreover, for both directionally unbiased regimes with *n* = 1 and *n* = 2.5, *AV G* (*t*) remains unchanged at 51 (see Fig. 2A1), while it increases linearly for the anterograde-biased regime with *n* = 1 and nonlinearly for the anterograde-biased regime with *n* = 2.5 (see Fig. 2B1). Notice that the diffusion coefficient *σ* is simply a time-scaling factor.

**Figure 2:**
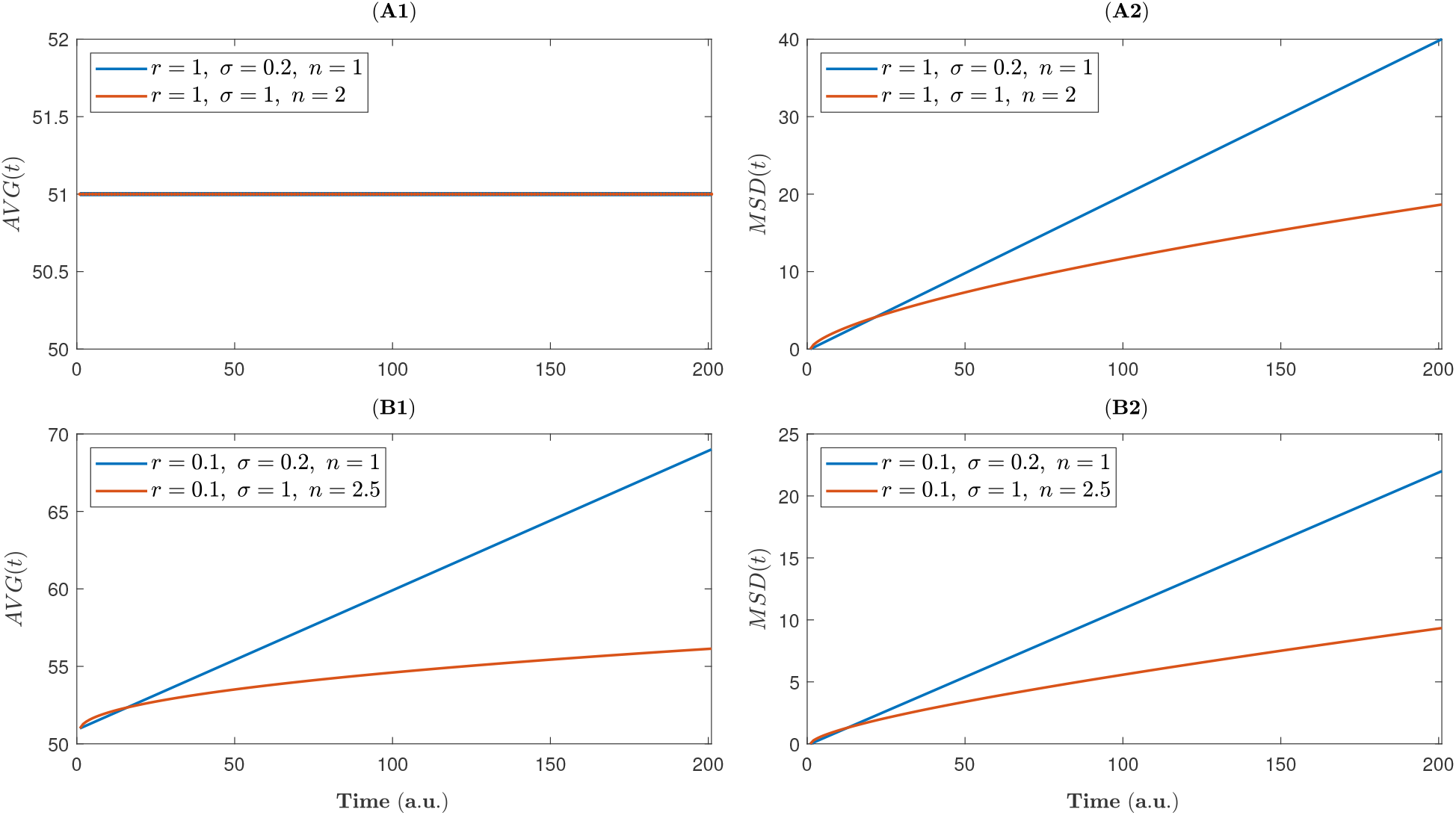
(A1) and (A2) present the average position *AV G* (*t*) and the mean squared displacement with respect to the average position *MSD* (*t*), respectively, for two directionally unbiased regimes (*r* = 1), and (B1) and (B2) also show *AV G* (*t*) and *MSD* (*t*), respectively, for two anterograde-biased regimes with *r* = 0.1 when the initial concentration at all 101 nodes was set to zero except node 51, which was set to 1.

## 7. Modeling tauopathy progression in the mouse brain

First discovered in 1975, tau is a multifunctional microtubule-associated protein (MAP) in the neuron, which many researchers have extensively studied its function to stabilize microtubules and encourage axonal prolongation [82, 83]. Tau protein is natively unfolded, and in physiological conditions its tendency for aggregation is low. However, there are modifications, such as phosphorylation and truncation [82], which may enable monomeric tau proteins to make aggregates. Tau aggregation characterizes neurodegenerative diseases known as tauopathies, including Alzheimer’s disease (AD), Huntington disease (HD), Pick disease (PiD), progressive supranuclear palsy (PSP), argyrophilic grain disease (AGD), corticobasal degeneration (CBD), and frontotemporal dementia with parkinsonism-17 (FTDP-17). Although the potential of tau protein to induce such diseases has been confirmed by the identification of tau mutants in patients with FTDP-17 [84], the mechanisms and pathways by which tau protein forms aggregates in tauopathies are not sufficiently comprehended.

Tau pathology induced by the injection of tau seeds into the mouse brain supports the idea that the harmful species can be transmitted from the inoculation sites to synaptically connected brain regions [85, 86]. Actually, the spreading of toxic tau species is chiefly attributed to the axonal transportation [87], and thus, due to the porous structure of axonal bundles [88–90], it is expected that, at the mesoscale, tau pathology is propagated by a nonlinear diffusion process. Further, cell culture and animal model studies have indicated that misfolded tau species can be transmitted trans-synaptically among neurons, both anterogradely and retrogradely [91, 92], and recently, researchers have also investigated directionally biased spreading of tauopathies using an in-silico model mimicking the two-neuron system [93] and a diffusion equation with the linear Laplacian operator on the mouse connectome [94]. This section proposes a model that captures the effects of both nonlinearity and directionality of spreading on the spatiotemporal evolution of tau pathology.

### 7.1. Constructing the Laplacian matrix using the Allen Mouse Brain Connectome

Here, we aim to take advantage of the publicly available data from the Allen Mouse Brain Connectivity Atlas (AMBCA) [95] (connectivity.brain-map.org) to construct a Laplacian matrix that meets our objectives. The inter-region connectivity matrix 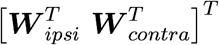 of the Allen Mouse Brain Connectome provides strength estimates of the mutual connections from 213 regions of interest (ROIs) in the right hemisphere to 426 (2 × 213) ROIs in the right (ipsilateral) and left (contralateral) hemispheres. Since the strength values resulted from a linear regression, a p-value was also assigned to each connection [95]. Fig. 3A. depicts this matrix for the maximum accepted p-value (the significance level) of 0.5.

**Figure 3:**
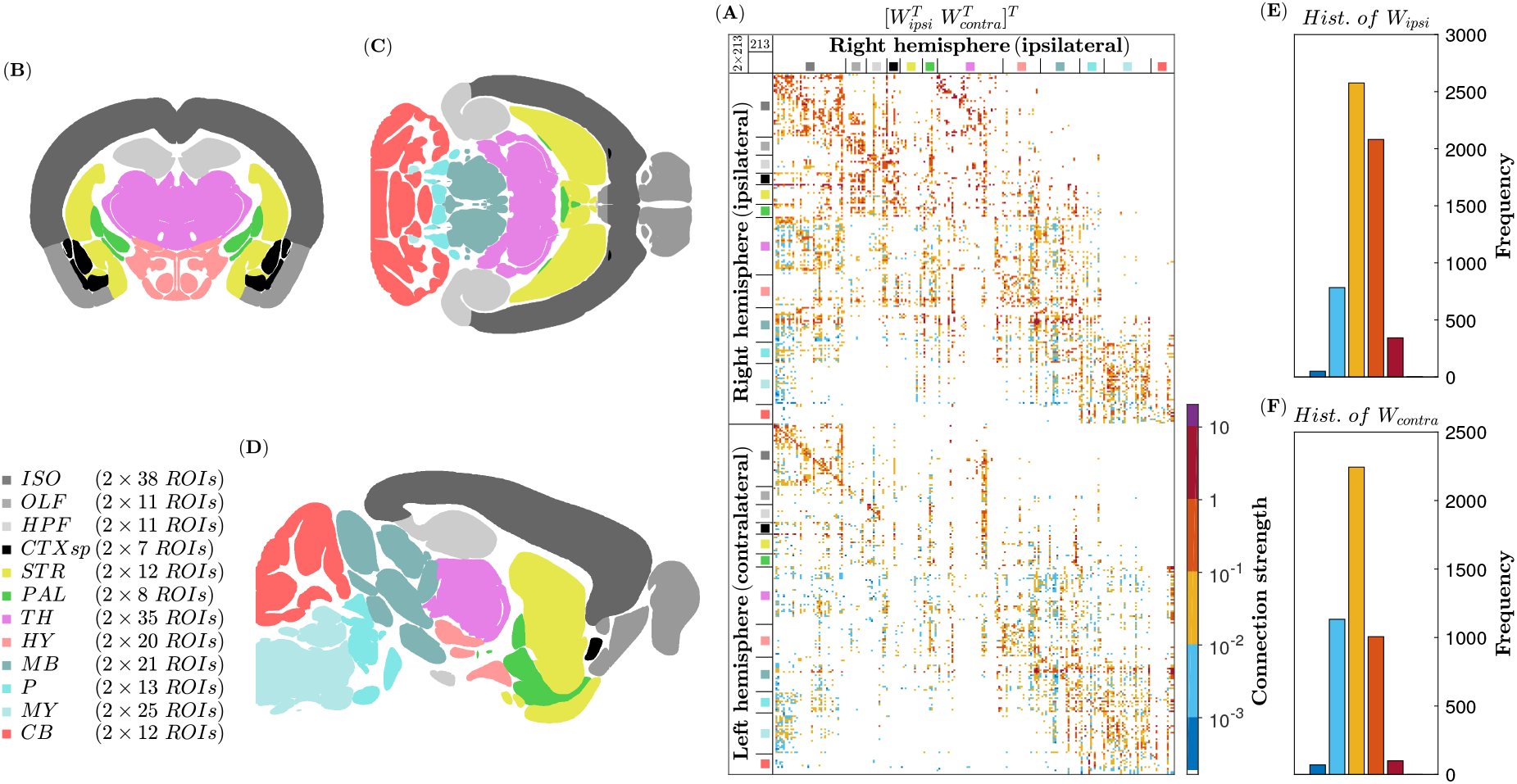
(A) Graphical representation of 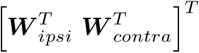 for the significance level of 0.5. Images (B), (C), and (D) show coronal, axial, and sagittal views of the mouse brain, respectively, which demonstrate the 12 major regions of the gray matter: isocortex (ISO), olfactory areas (OLF), hippocampal formation (HPF), cortical subplate (CTXsp), striatum (STR), pallidum (PAL), thalamus (TH), hypothalamus (HY), midbrain (MB), pons (P), medulla (MY), and cerebellum (CB). To generate this result, we used the dataset of the Allen Mouse Common Coordinate Framework (CCFv3) [96] (help.brain-map.org/display/mouseconnectivity/API, specifically annotation/ccf 2017). (E) and (F) also display the histograms of ***W** _ipsi_* and ***W** _contra_*, respectively.

In the AMBCA project, axonal projections were traced by injecting recombinant adeno-associated virus (AAV) expressing enhanced green fluorescent protein (EGFP), which is known as an anterograde tracer. In this sense, based on the assumption that hemispheric symmetry holds, one can define an anterograde weighted adjacency matrix in the following way:

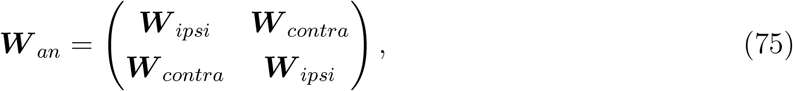

and, consequently, it makes sense to suggest a retrograde weighted adjacency matrix 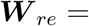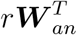 with *r* ∈ ℝ_>_0. The two weighted adjacency matrices may eventually be combined into a single matrix

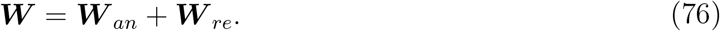

Now, let us construct the adjacency matrix ***A*** by replacing the nonzero elements of ***W*** with 1. Using mathematical induction, it can be shown that the number of different paths of length *k* from node *i* to node *j*, where *k* is a positive integer, equals to (*j, i*)-th entry of ***A**^k^*. Thus, there exists at least one path of length lower than or equal to *p\** from each node to every other node if the matrix 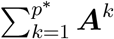 has positive values in all its off-diagonal entries. The minimum value of *p\** for different significance levels are illustrated in Table 1, which ensures that the weighted connectivity matrix ***W*** represents a strongly connected symmetric digraph for significance levels greater than or equal to 10^−5^.

**Table 1:**
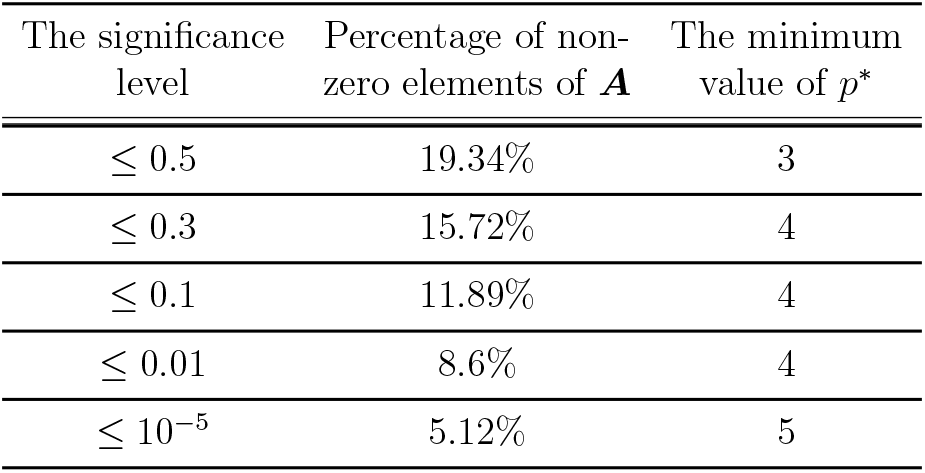
Percentage of nonzero elements of the adjacency matrix ***A*** and the corresponding minimum value *p\** for different maximum accepted p-values of the connection strengths.

Normalizing the matrix ***W*** requires taking into account the fact that the resulting Laplacian matrix must satisfy the condition **1***^T^ **L*** = 0, ensuring that the total concentration will be preserved during a diffusion process. Thus, one may propose two methods to normalize ***W*** : either by converting ***W*** into a left stochastic matrix, i.e., a matrix with each column summing to 1, or by dividing ***W*** by ‖***W*** ‖_1_ where ‖.‖_1_ denotes the maximum absolute column sum of the matrix. It is more preferable to go with the latter option, which is simply scaling ***W*** by a coefficient, as the ratio of any two elements from different columns of ***W*** remains unchanged after normalizing. Moreover, since ‖***W*** ‖1 depends on *r*, 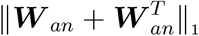 will be substituted here for ‖***W*** ‖_1_ to ensure that the scaling coefficient is continuously differentiable with respect to *r*, which is necessary to calculate the sensitivity function that will be discussed in Subsection 7.3. Lastly, the normalized weighted adjacency matrix

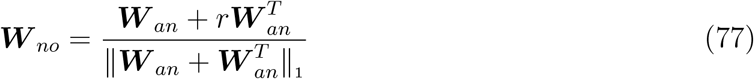

yields the Laplacian matrix

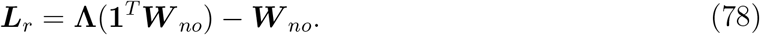

### 7.2. Simulation results

Taking into account the Laplacian matrix ***L**_r_*, let us rewrite Eq. (45) for the mouse connectome *G* as follows:

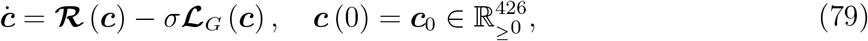

where ***α*** = *α***1**, ***ρ*** = *ρ***1**, ***μ*** = *μ***1**, ***ν*** = *ν***1**, ***n*** = *n***1**, and the parameter vector ***θ*** = (*α, ρ, μ, ν, σ, r, n*)^*T*^ belongs to the set

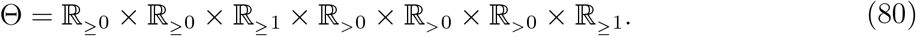

Having analyzed the data from transgenic tau P301S mice, which represent animal models of tauopathy, for up to 4 months, Meisl et al. [97] reported a doubling time of approximately 2 weeks in the brain stem, neocortex, and frontal lobe. This gives us a crude estimate of *α*, which is *ln*(2)/2 ≈ 0.35. In animal experiments examining tauopathy progression, tau prion strains are commonly injected into the ROIs of the hippocampal formation (HPF) such as *CA*1 [98]. In our simulations, we therefore consider the following initial concentrations:

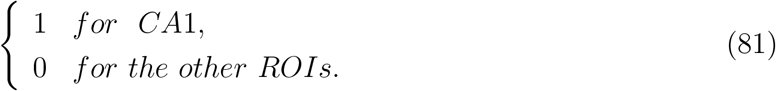

We conducted two in-silico experiments to examine the effect of nonlinear diffusion on the spatiotemporal evolution of tauopathy. First, a simulation was carried out for an anterograde-biased regime with *r* = 0.1 and the initial concentrations specified in Eq. (81), where *α* = 0.35, and all the other parameters were set to 1; See Fig. 4. The simulation was then repeated with the same initial conditions and parameters, except that *n* was changed to 1.5. Fig. 5 illustrates how tauopathy evolved in the second in-silico experiment relative to that in the first. As shown in Fig. 5, increasing *n* slowed down the spreading of tau pathology. In fact, it resulted in tauopathy progression being localized more significantly to the ROIs adjacent to the in-silico injection site *CA*1 during the simulation period. One may note that decreasing the diffusion coefficient *σ* can also lead to a similar result. To address this claim and investigate the effect of parameter variations on the progression of tauopathy, we intend to perform a sensitivity analysis.

**Figure 4:**
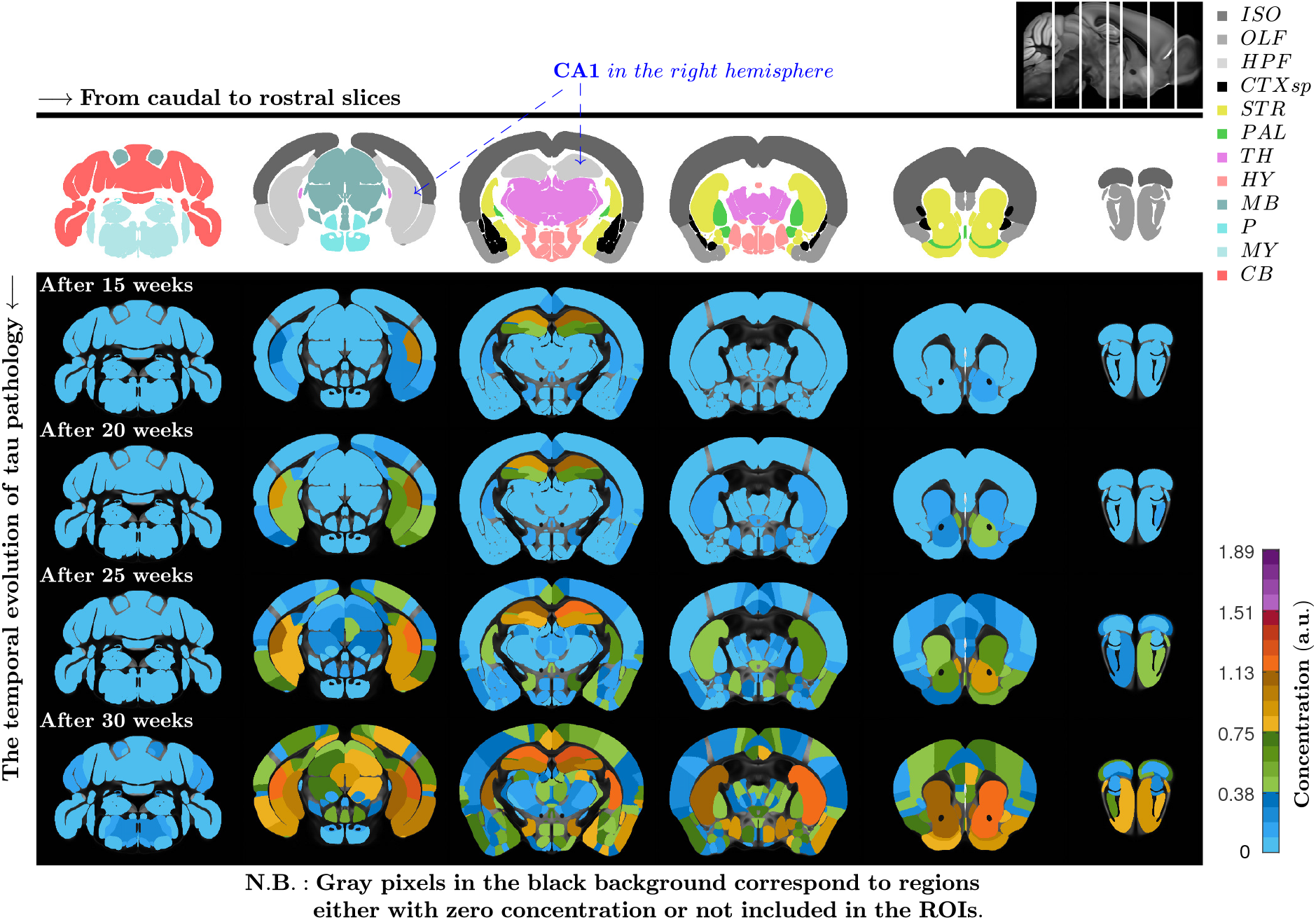
It displays the in-silico spatiotemporal evolution of tau pathology for the initial concentrations given in Eq. (81) when (*α, ρ, μ, ν, σ, r, n*) = (0.35, 1, 1, 1, 0.1, 1). To generate this result, we used the dataset of the Allen Mouse Common Coordinate Framework (CCFv3) [96] (help.brain-map.org/display/mouseconnectivity/API, specifically annotation/ccf 2017).

**Figure 5:**
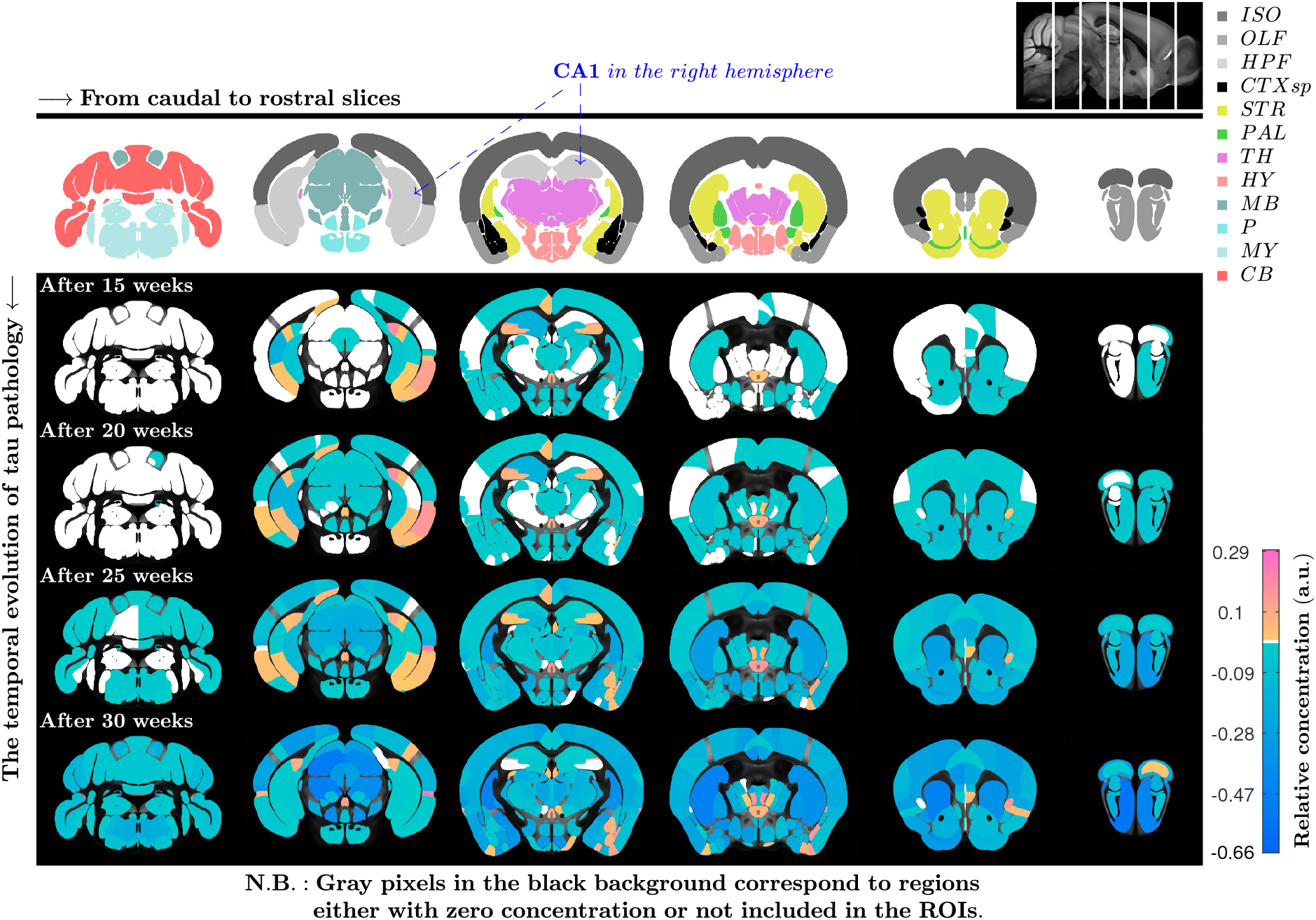
It exhibits the relative spatiotemporal evolution of tauopathy for the initial concentrations given in Eq. (81) in the in-silico experiment with the parameters (*α, ρ, μ, ν, σ, r, n*) = (0.35, 1, 1, 1, 0.1, 1.5) in comparison with that of one with the same parameters, except that *n* = 1. To generate this result, we used the dataset of the Allen Mouse Common Coordinate Framework (CCFv3) [96] (help.brain-map.org/display/mouseconnectivity/API, specifically annotation/ccf 2017).

### 7.3. Sensitivity analysis

Let 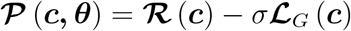 ; then, by integration, the system (79) can be written as

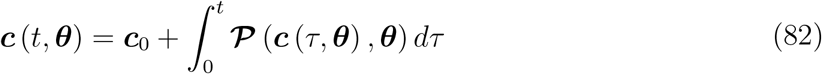

having a unique solution ***c*** (*t, **θ***) for an initial concentration 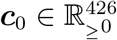 and a parameter vector ***θ*** ∈ Θ by Corollary 5.1. Calculating the partial derivatives of 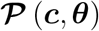 with respect to ***c*** and ***θ*** yields

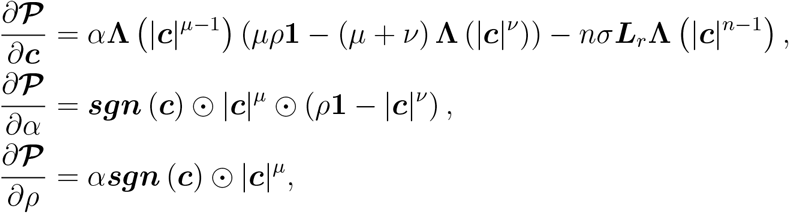

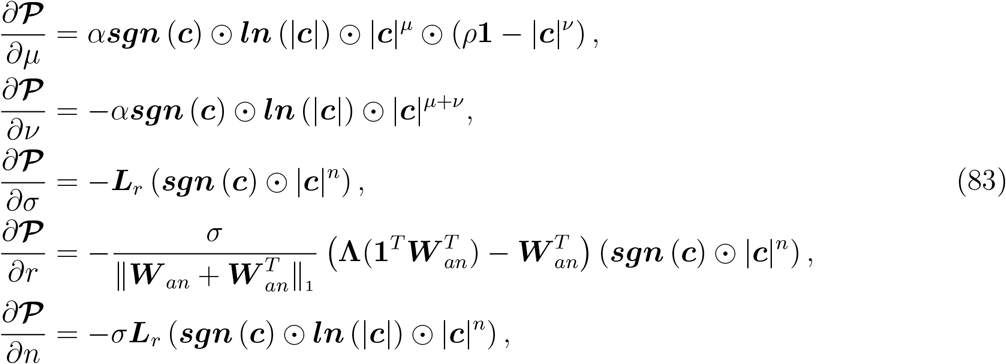

where are continuous for all (***c**, **θ***) ∈ ℝ^426^ × Θ^*\**^ with

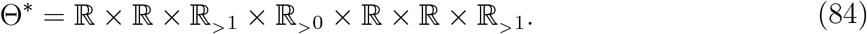

Note that lim_***c**→*0_ ***ln*** (|***c***|) ⊙ |***c***|^*n*^ = 0 for *n* > 0. Thanks to the continuous differentiability of 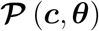 with respect to ***c*** and ***θ***, the solution ***c*** (*t, **θ***) is differentiable with respect to ***θ*** near a nominal value ***θ***_0_ ∈ Θ^*\**^. Thus, we can obtain the following equation by taking partial derivatives of Eq. (82) with respect to ***θ***.

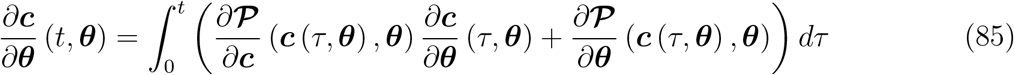

For a nominal solution ***c*** (*t, **θ***_0_) with ***θ***_0_ ∈ Θ ∩ Θ^*\**^, differentiating both sides of Eq. (85) with respect to *t* gives the sensitivity equation

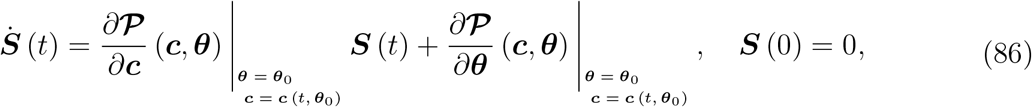

where 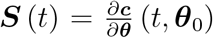. The sensitivity function ***S*** (*t*) provides a first-order estimate of the effect of parameter variations on the solution ***c*** (*t, **θ***_0_). In other words, for ***θ*** sufficiently close to ***θ***_0_, the solution ***c*** (*t, **θ***) can be approximated by a Taylor series about the solution ***c*** (*t, **θ***_0_), i.e.,

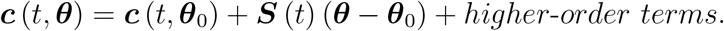

Due to the large number of elements in the sensitivity function ***S*** (*t*), i.e., 426 × 7, it is difficult to scrutinize the simulation results for this function, and therefore a version of ***S*** (*t*) with a lower dimension is required. For the solution ***c*** (*t, **θ***), let us now define the spatial average concentration at time *t*

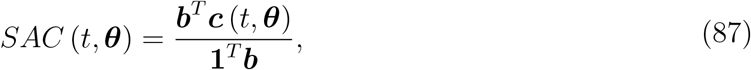

where the *i*-th element of the vector ***b*** is the number of voxels occupied by the *i*-th ROI, according to the dataset of the Allen Mouse Common Coordinate Framework (CCFv3) [96] (help.brain-map.org/display/mouseconnectivity/API, specifically annotation/ccf 2017). The spatial average concentration approximately represents the spatial average of the distribution of the species concentration over the entire brain. It is easy to see that the sensitivity function for the spatial average concentration is derived as follows:

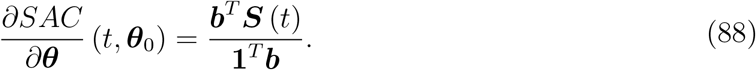

Taking into account the initial concentrations given in Eq. (81), we computed the sensitivity function ***S*** (*t*) and its corresponding function *∂SAC/∂**θ*** (*t, **θ***_0_) at 16 different points ***θ***_0_ in the parameter space. The maximum absolute value over time of each element of *∂SAC/∂**θ*** (*t, **θ***_0_) have been reported in Table 2 for the chosen parameter values. Fig. 6 also illustrates the simulation results from which we derived the maximum absolute values in the first two rows of Table 2. As outlined in Table 2, the spatial average concentration of tauopathy generally appears to be more sensitive to variations in the parameter *μ* than to variations in the other parameters. Among the reaction term coefficients, including *α*, *ρ*, *μ*, and *ν*, the variation in *ν* also has the least impact on the model output at the selected points in the parameter space, and in general, there is a greater sensitivity to variations in *α* than that to variations in *ρ*. However, as indicated in Fig. 6, variations in *ρ* can substantially change the steady state of the spatial average concentration. According to Table 2, when restricting ourselves to the diffusion term coefficients, namely, *σ*, *r*, and *n*, the average concentration of tau pathology seems chiefly to be most sensitive to *r* at the points in the parameter space with an anterograde-biased regime (*r* = 0.1) and to *n* at the points with a directionally unbiased regime (*r* = 1). Additionally, among all the seven parameters considered, variations in the parameter *σ* have the least influence on the model output.

**Table 2:**
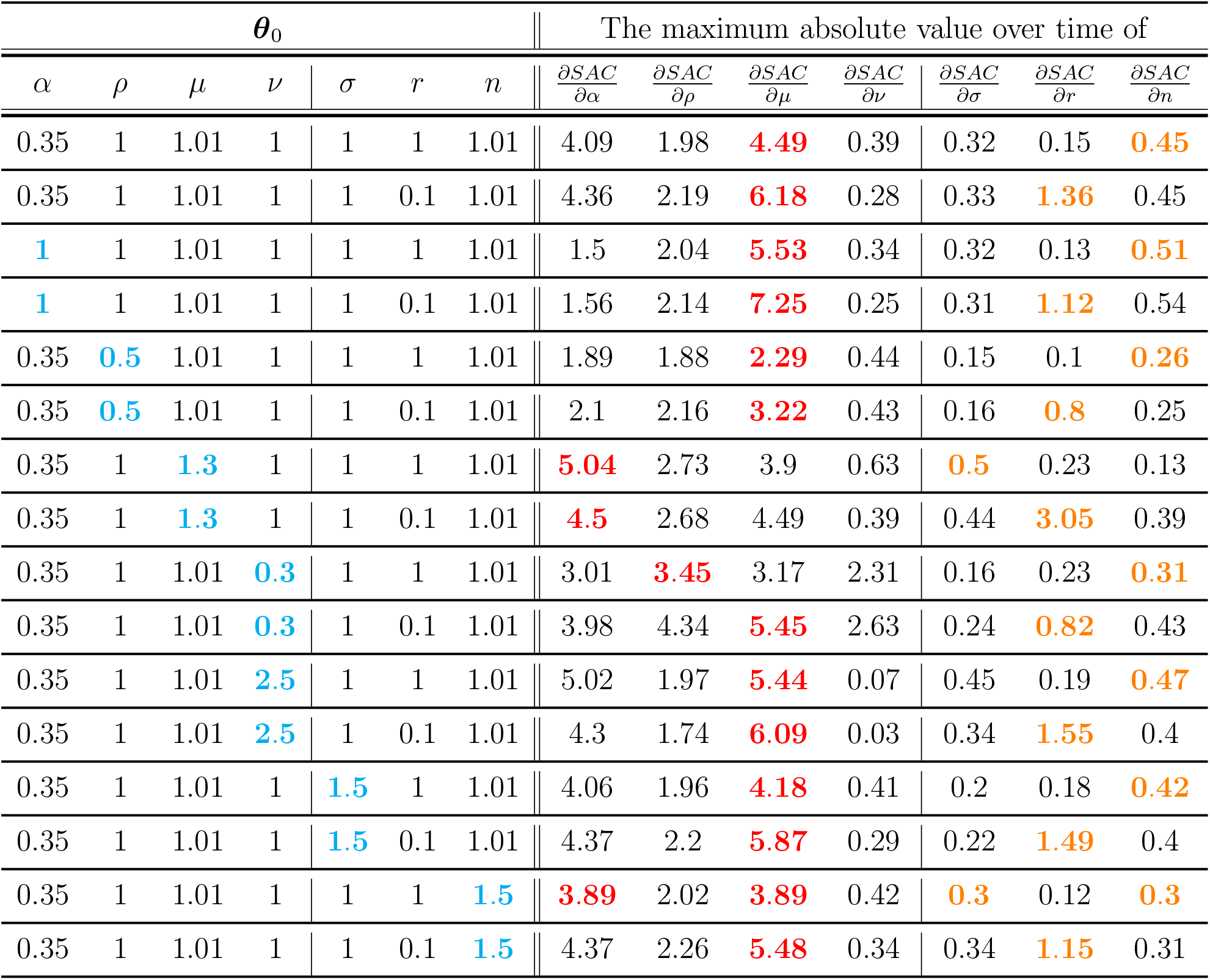
The maximum absolute value over time of the elements of *∂SAC/∂**θ*** (*t, **θ***_0_) for some different nominal values ***θ***_0_, where the initial concentrations are taken as Eq. (81).

**Figure 6:**
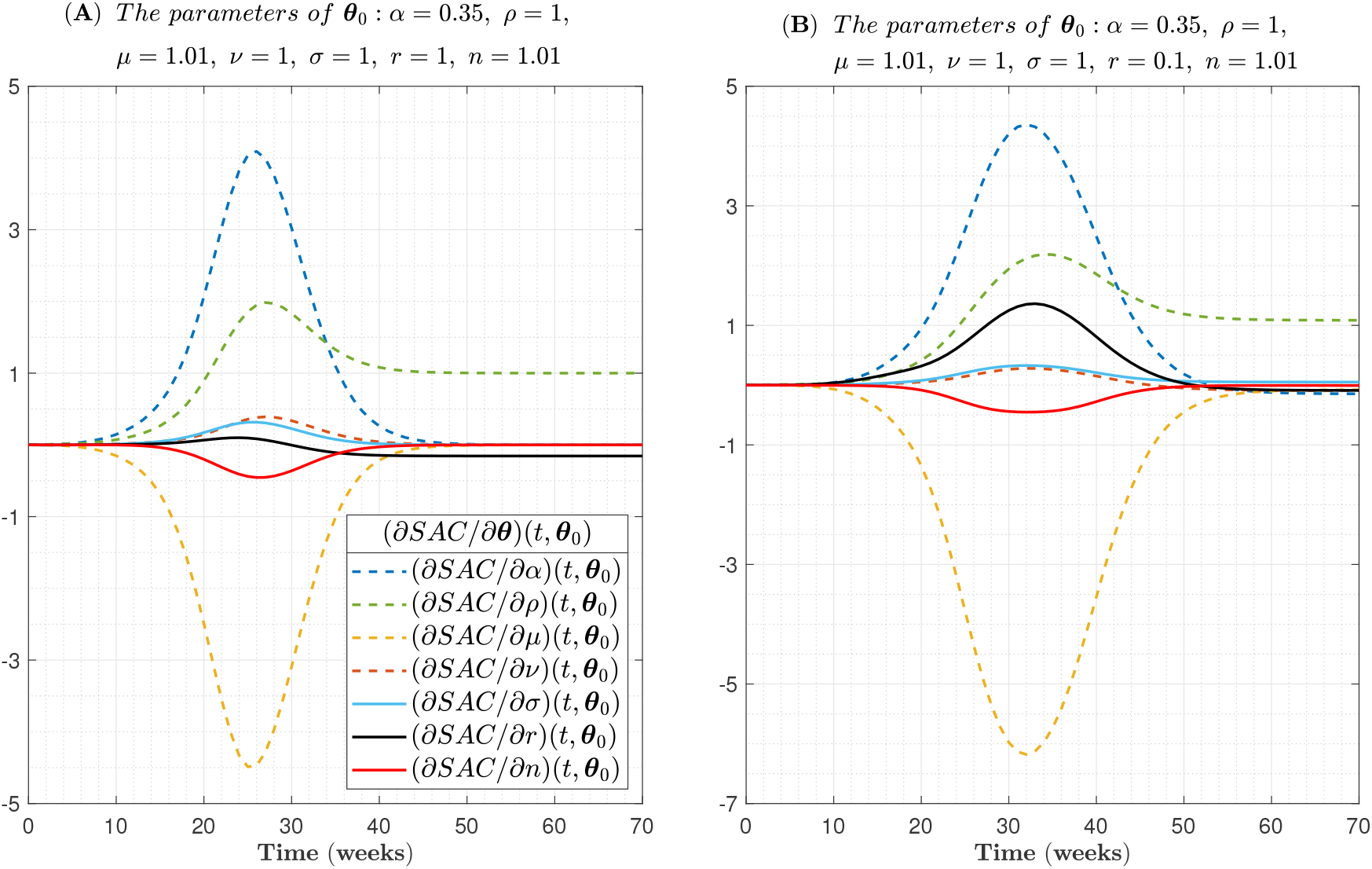
The elements of *∂SAC/∂**θ*** (*t, **θ***_0_) are depicted in (A) for a directionally unbiased regime and in (B) for an anterograde-biased regime with r=0.1, where the initial concentrations are taken as Eq. (81).

## 8. Conclusion

In this manuscript, the extended FP diffusion and Fisher-KPP reaction-diffusion equations on weighted digraphs whose SCCs are terminals were introduced. For the extended FP diffusion equation, it was shown that all solutions remain bounded and converge to equilibria which are stable. Further, we found positively invariant sets containing exactly one equilibrium point. Indeed, the domain of attraction of each equilibrium point was obtained. We also presented a family of positively invariant sets for the extended FP Fisher-KPP reaction-diffusion equation. Since a weighted undirected graph can be considered as a symmetric digraph, our results are also applicable to any arbitrary weighted undirected graph. To assess the practical significance of the proposed equations, it was first illustrated that the extended FP diffusion equation can generate a directionally-biased anomalous subdiffusion process on a one-dimensional lattice, which can be applied to describe the motion of motor proteins on microtubules. Moreover, we modeled tauopathy progression in the mouse brain connectome using the extended FP Fisher-KPP equation and then conducted a sensitivity analysis to estimate the effect of model parameter variations on solutions.

Here, the active concentration was also introduced. By definition, the active concentration of a given species can take a negative value, indicating the dominance of an antibody of that species. The concept of active concentration combined with the proposed extended FP equations enables us to capture the dynamics of a given species in presence of its antibody, which provides a useful framework for future research on studying antibodies, such as anti-tau and anti-amyloid antibodies for Alzheimer’s disease.

## Data availability

In this research project, we drew on the publicly available data from the Allen Mouse Brain Connectivity Atlas (AMBCA) [95] (connectivity.brain-map.org) and the dataset of the Allen Mouse Common Coordinate Framework (CCFv3) [96] (help.brain-map.org/display/mouseconnectivity/API, specifically annotation/ccf 2017). All computations, including numerical solution of differential equations, were also done using MATLAB.

## Acknowledgments

We acknowledge the support of the Natural Sciences and Engineering Research Council of Canada (NSERC) and GCS Faculty Research Support, Concordia University.

